# Functional Characterization of *lncRNA152* as an Angiogenesis-Inhibiting Tumor Suppressor in Triple-Negative Breast Cancers

**DOI:** 10.1101/2022.02.09.479778

**Authors:** Dae-Seok Kim, Shrikanth S. Gadad, Rohit Setlem, Kangsan Kim, Srinivas Malladi, Tim Y. Hou, Tulip Nandu, Cristel V. Camacho, W. Lee Kraus

**Author notes:** <u>Lead Contact (for manuscript correspondence)</u>: W. Lee Kraus, Ph.D., Cecil H. and Ida Green Center for Reproductive Biology Sciences, The University of Texas Southwestern Medical Center at Dallas, 5323 Harry Hines Boulevard, Dallas, TX 75390-8511. **<u>Disclosures</u>**: The authors have no competing interests to declare for this work.

## Abstract

Long non-coding RNAs have been implicated in many of the hallmarks of cancer. Herein, we found that the expression of *lncRNA152* (*lnc152*; a.k.a. *DRAIC*), which we annotated previously, is highly upregulated in luminal breast cancer (LBC) and downregulated in triple-negative breast cancer (TNBC). Knockdown of *lnc152* promotes cell migration and invasion in LBC cell lines. In contrast, ectopic expression of *lnc152* inhibits growth, migration, invasion, and angiogenesis in TNBC cell lines. In mice, *lnc152* inhibited the growth of TNBC cell xenografts, as well as metastasis of TNBC cells in an intracardiac injection model. Transcriptome analysis of the xenografts indicated that *lnc152* downregulates genes controlling angiogenesis. Using pull down assays coupled with LC-MS/MS, we identified RBM47, a known tumor suppressor in breast cancer, as a *lnc152*-interacting protein. The effects of *lnc152* in TNBC cells are mediated, in part, by regulating the expression of RBM47. Collectively, our results demonstrate that *lnc152* is an angiogenesis-inhibiting tumor suppressor that attenuates the aggressive cancer-related phenotypes found in TNBC.

**Statement of Significance:** This study identifies *lncRNA152* as an angiogenesis-inhibiting tumor suppressor that attenuates the aggressive cancer-related phenotypes found in TNBC by upregulating the expression of the tumor suppressor RBM47. As such, *lncRNA152* may serve as a biomarker to track aggressiveness of breast cancer, as well as therapeutic target for treating TNBC.

## Introduction

Metastasis leads to the spread of cancer from its primary site of origin to a distant site via the circulatory system to establish a secondary tumor. The process begins with invasion and intravasation into the surrounding lymphatics and blood vessels, and culminates with colonization of the disseminated tumor cells in a distal organ, angiogenesis, and growth (1,2). Angiogenesis is an essential component of the metastatic pathway. New blood vessels promote tumor metastasis by providing the route of exit for tumor cells from the primary tumor and colony formation at secondary sites (3,4). Moreover, new blood vessels within the tumor are needed to provide sufficient oxygen and nutrients to support tumor growth (3,4). As such, anti-angiogenic therapies have gained significant attention as a potential therapeutic approach for cancer treatment (5,6). In breast cancers, metastasis is the major cause of breast cancer-related morbidity and mortality. Breast cancer preferentially metastasizes to the lungs, brain, bone, liver, and distal lymph-nodes (7,8). A better understanding of the molecular mechanisms regulating metastasis is crucial for development of novel therapeutic strategies.

Long non-coding RNAs (lncRNAs) are non-protein coding RNA transcripts with lengths greater than 200 nucleotides, with many thousands of lncRNA genes identified in the human genome (9,10). LncRNAs mediate a host of disparate molecular and cellular functions, including gene regulation (9,10). Increasingly, lncRNAs have been implicated in many of the hallmarks of cancer, such as proliferation, growth suppression, motility, immortality, angiogenesis, and viability (11,12). Numerous lncRNAs have been found to be mutated or abnormally expressed in various cancer types (13,14), suggesting that they can potentially be used as prognostic biomarkers and therapeutic targets in cancer.

The biological functions of lncRNAs in cancer are diverse, with connections to many different aspects of cancer biology. LncRNAs may function as tumor suppressors or oncogenes during cancer progression (15). Increasing evidence indicates that lncRNAs can mediate tumor suppressive functions in a wide variety of prevalent cancer types (15-19). LncRNAs have also been shown to play a role in metastasis-related functions, such as angiogenesis (10,15). LncRNAs play a critical role in tumor angiogenesis by regulating the various underlying biological processes that control angiogenesis, including epigenetic mechanisms that control angiogenic gene expression programs (20,21). The diversity of lncRNA expression, sequences, structures, interaction partners, subcellular localization, and functions makes them ideally suited to control a broad array of critical cancer-related biological functions.

In a previous study, we used genomics and bioinformatics to discover and annotate lncRNAs controlling cell cycle gene expression and proliferation in breast cancer cells, including *lncRNA152* (*lnc152*; a.k.a. *DRAIC*), which is located in a tumor suppressor locus on human chromosome 15q23 (22). In a series of functional assays, we observed key roles for *lnc152* in proliferation, cell cycle progression, and regulation of the estrogen signaling pathway in breast cancer cells (22). A contemporaneous study identified *lnc152/DRAIC* as a tumor suppressor in prostate cancer (23). Subsequent studies have affirmed a key role for *lnc152/DRAIC* in the biology of an array of cancers (24-30), but the precise mechanisms by which it suppresses a wide range of cancer-related behaviors, such as proliferation, migration, invasion, metastasis, and angiogenesis, remain largely unknown. Here, our studies demonstrate that *lnc152* exhibits elevated expression in the luminal subtype of breast cancer, but is down-regulated in the triple-negative (TN)/basal subtype of breast cancer. *Lnc152* inhibits cancer-related phenotypes, including growth, migration, invasion, metastasis, and angiogenesis by upregulating the expression of the RNA-binding tumor suppressor protein, RBM47, to control a tumor suppressing gene expression program.

## Results

### *Lnc152* is highly expressed in luminal breast cancer and acts to attenuate cell migration and invasion

We previously identified *lnc152* in MCF-7 luminal human breast cancer cells (22) (Figure S1, A and B). To further functionally characterize *lnc152* and its role in breast cancer, we examined its expression in breast cancer patient samples and cell lines. We observed that *lnc152* is highly expressed in breast cancers compared to normal breast tissues using TCGA data (Figure 1A). Moreover, we found that *lnc152* is upregulated in luminal and HER2 breast cancers, but downregulated in basal-like breast cancer (Figure 1B). Similar results were observed in breast cancer cell lines (Figure 1C). In perturbation-response experiments, siRNA-mediated knockdown of *lnc152* promoted the migration of and invasion by MCF-7 cells in culture-based assays (Figure 1, D and E). siRNA-mediated knockdown of *lnc152* also altered gene expression in MCF-7 cells as assessed by RNA-seq, including the upregulation of many genes involved in cancer-related biological processes, such as extracellular matrix organization, and cell migration (Figure 1F).

**Figure 1.**
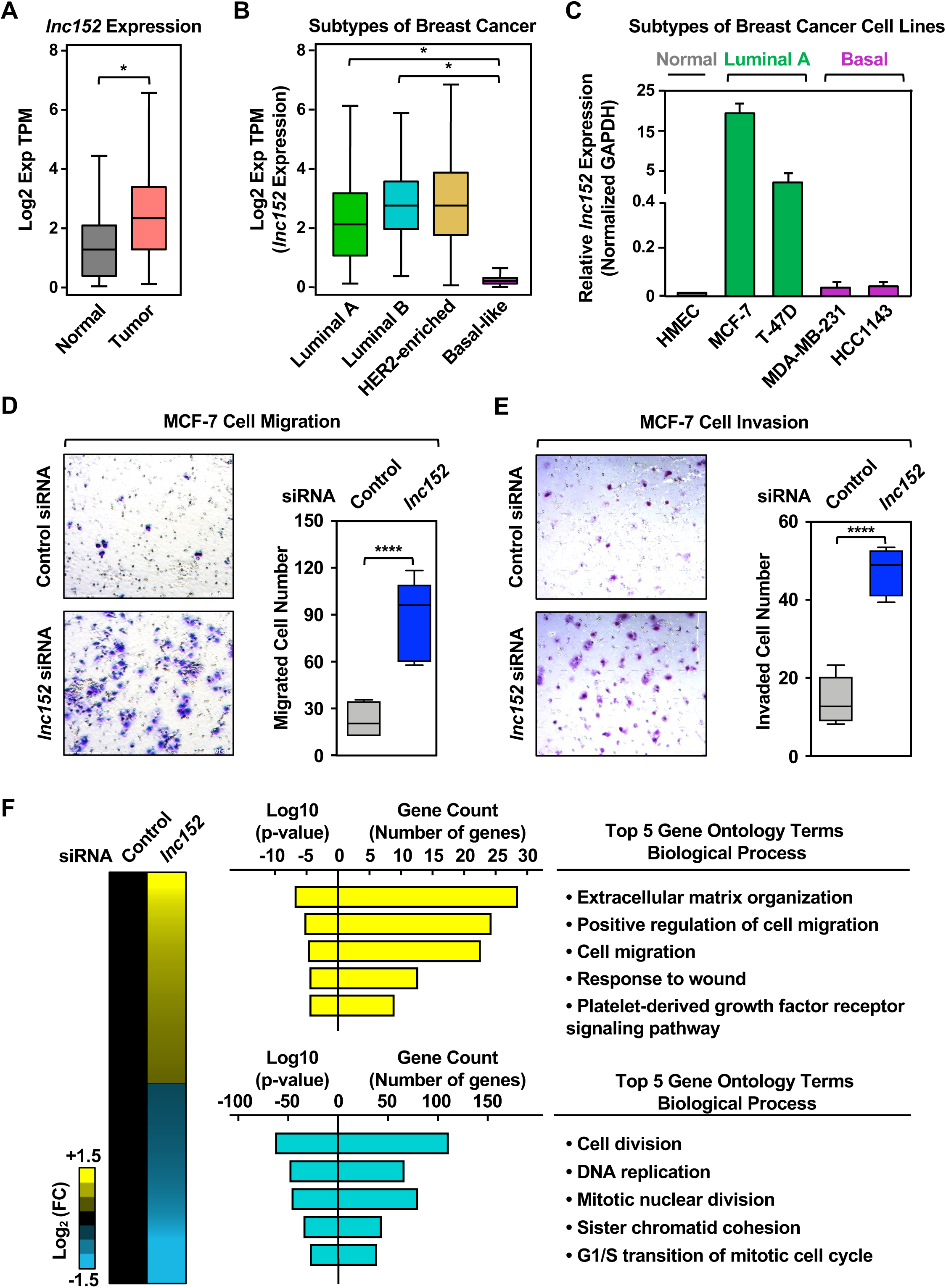
*Lnc152* is differentially expressed across the distinct molecular subtypes of breast cancer and is required for suppression of luminal breast cancer cell invasiveness. **(A)** RNA-seq expression data for *lnc152* in primary tumors (TCGA breast cancer samples, n = 1085) compared to normal breast tissues (TCGA normal and GTEx data, n = 291). Bars marked with asterisks are statistically different from each other (two-tailed Student’s t test, * p < 0.05). **(B)** Box plots comparing the expression of *lnc152* in different molecular subtypes of breast cancers [luminal A (n = 415), luminal B (n = 194), HER2-enriched (n = 66), and basal-like (n = 135)]. Bars marked with asterisks are statistically different from each other (two-tailed Student’s t test, * p < 0.05). **(C)** Bar graphs showing the expression of *lnc152* in breast cancer cell lines representing different molecular subtypes. **(D)** (*Left*) siRNA-mediated knockdown of *lnc152* promotes the cell migration of MCF-7 cells (luminal A breast cancer subtype) compared to knockdown with a control siRNA. (*Right*) Quantification of the results from the experiments shown in the left panel. Each box plot represents the mean ± SEM, n = 7. Bars marked with asterisks are statistically different from each other (two-tailed Student’s t test, **** p < 0.0001). **(E)** (*Left*) siRNA-mediated knockdown of *lnc152* promotes the cell invasion of MCF-7 cells (luminal A breast cancer subtype) compared to knockdown with a control siRNA. (*Right*) Quantification of the results from the experiments shown in the left panel. Each box plot represents the mean ± SEM, n = 6. Bars marked with asterisks are statistically different from each other (two-tailed Student’s t test, **** p < 0.0001). **(F)** siRNA-mediated knockdown of *lnc152* alters gene expression in MCF-7 cells. (*Left*) Heatmap showing the results of RNA-seq assays from *lnc152*-depleted MCF-7 cells. (*Middle*) Gene ontology analysis showing biological processes that are enriched (yellow) and those that are de-enriched (cyan). (*Right*) GO terms in each cluster indicating the top five biological processes determined by REViGO analysis.

The enrichment and function of *lnc152* in the luminal breast cancer subtype is reflected in its regulation by FoxA1, a key transcription factor in luminal breast cancers (31-34) (Figure S1C). In this regard: (1) FoxA1 binds at the promoter and upstream regions of the *lnc152* gene (Figure S1B), (2) the expression of *FOXA1* mRNA and *lnc152* are correlated in public gene expression data sets (Figure S1D-S1F) and luminal breast cancer cell lines (Figure S1G). Finally, siRNA-mediated knockdown of FoxA1 decreases the expression level of *lnc152* in luminal A breast cancer cell lines (Figure S1H). Collectively, these results indicate that *lnc152* is highly expressed in luminal breast cancers and downregulated in triple-negative/basal breast cancers. In addition, we demonstrate that *lnc152* regulates a gene expression program that promotes cell migration and invasion by luminal breast cancer cells.

### Ectopic expression of *lnc152* inhibits cancer-related cellular phenotypes in triple-negative breast cancer cells, including angiogenesis

As shown above, *lnc152* is significantly downregulated in basal-like breast cancer and triple-negative breast cancer (TNBC) cells, suggesting that the low levels of *lnc152* may be associated with more aggressive cancer features. To determine whether *lnc152* can modulate the aggressiveness of TNBC cells, we assayed a range of cancer-related cellular phenotypes, such as cell proliferation, migration, invasion, and blood vessel formation, using TNBC cells (i.e., MDA-MB-231 and HCC1143) engineered for Dox-inducible ectopic expression of *lnc152*, or GFP mRNA as a control (Figure 2A, Figure S2A). Dox-induced ectopic expression of *lnc152* inhibited the proliferation of MDA-MB-231 cells (Figure 2B), but not HCC1143 cells (Figure S2B), compared to ectopic expression of GFP mRNA (Figure 2B, Figure S2B). In contrast, Dox-induced ectopic expression of *lnc152* inhibited migration and invasion for both cell lines, compared to ectopic expression of GFP mRNA (Figure 2, C and D; Figure S2, C and D). While both HCC1143 and MDA-MB-231 are TNBC cell lines, HCC1143 (derived from ductal carcinoma) and MDA-MB-231 (derived from adenocarcinoma) can be further stratified into different classes with different phenotypes (35), which may account for the different effects of *lnc152* expression in these two cell lines.

**Figure 2.**
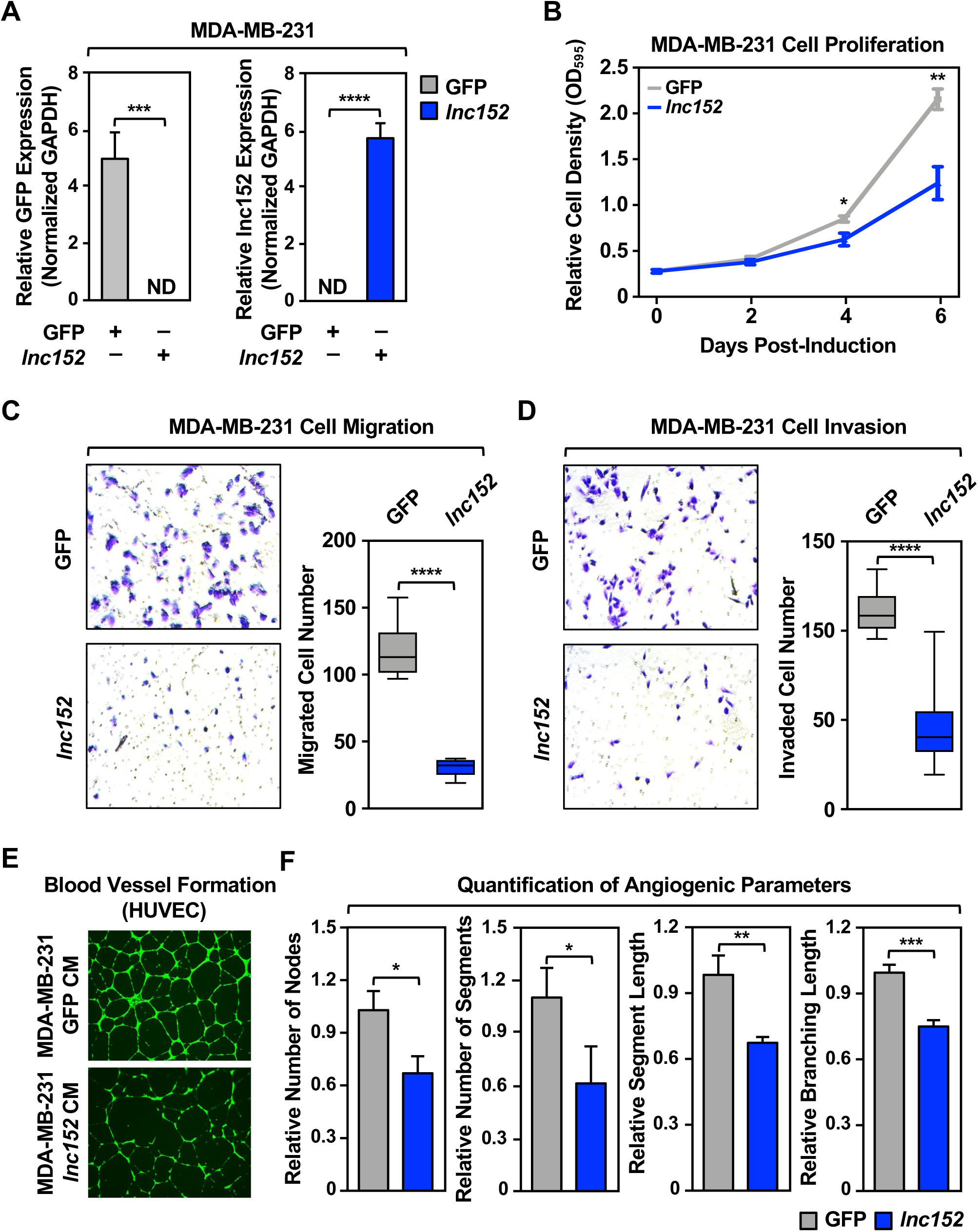
Ectopic expression of *lnc152* inhibits aggressive cancer-related outcomes in MDA-MB-231 TNBC cells. **(A)** Generation of MDA-MB-231 (basal breast cancer subtype) cell lines for Dox-inducible ectopic expression of GFP or *lnc152*. GFP mRNA and *lnc152* levels were determined by RT-qPCR and normalized to GAPDH mRNA. ND indicates expression level was not detectable. Each bar represents the mean + SEM, n = 3. Bars marked with asterisks are statistically different from each other (two-tailed Student’s t test, *** p < 0.001, **** p < 0.0001). **(B)** Dox-induced ectopic expression of *lnc152* inhibits MDA-MB-231 cell proliferation compared to ectopic expression of GFP mRNA. Cells were treated daily with 0.5 μg/mL of Dox for six days. Each point represents the mean ± SEM, n = 3. Points marked with asterisks are statistically different from each other (two-tailed Student’s t-test, * p < 0.05, ** p < 0.01). **(C)** (*Left*) Dox-induced ectopic expression of *lnc152* inhibits the migration of MDA-MB-231 cells compared to ectopic expression of GFP mRNA. (*Right*) Quantification of the results from the experiments shown in the left panel. Each bar represents the mean ± SEM, n = 8. Bars marked with asterisks are statistically different from each other (two-tailed Student’s t test, **** p < 0.0001). **(D)** (*Left*) Dox-induced ectopic expression of *lnc152* inhibits the invasion of MDA-MB-231 cells compared to ectopic expression of GFP mRNA. (*Right*) Quantification of the results from the experiments shown in the left panel. Each bar represents the mean ± SEM, n = 8. Bars marked with asterisks are statistically different from each other (two-tailed Student’s t test, **** p < 0.0001). **(E)** Dox-induced ectopic expression of *lnc152* prevents human umbilical vein endothelial cell (HUVEC) tube formation on Matrigel. Morphological appearance of HUVEC grown on Matrigel with conditioned medium (CM) collected from MDA-MB-231 cells expressing Dox-induced GFP or *lnc152*, stained with Calcein-AM (green) and detected by fluorescence microscopy. **(F)** Quantification of the results from the experiments shown in panel E. Image J software with the Angiogenesis plugin was used to detect the number of nodes, number of segments, segment length, and branching length. Each bar represents the mean + SEM, n = 4. Bars marked with asterisks are statistically different from each other (two-tailed Student’s t test, * p < 0.05, ** p < 0.01, *** p < 0.001).

Importantly, conditioned medium (CM) collected from MDA-MB-231 cells with Dox-induced expression of *lnc152*, but not GFP, prevented tube formation by human umbilical vein endothelial cells (HUVECs) on Matrigel (Figure 2E and 2F), an in vitro assay of angiogenesis (36). These results show that *lnc152* can inhibit cancer-related cellular phenotypes in triple-negative breast cancer cells. This includes inhibition of angiogenic phenotypes in HUVECs by TNBC cells whose proliferation is inhibited by *lnc152* expression (e.g., MDA-MB-231 cells, not HCC1143 cells) through production of a putative soluble factor.

### *Lnc152* acts as a tumor suppressor in TNBC cells

To investigate the inhibition of cancer-related phenotypes by *lnc152* in vivo, we monitored the growth and biology of xenograft tumors formed in nude mice from MDA-MB-231 cells engineered for Dox-inducible ectopic expression of *lnc152*. We observed that ectopic expression of *lnc152* caused a significant reduction in the growth of the xenograft tumors (Figure 3A-C). Moreover, ectopic expression of *lnc152* caused a significant reduction in blood vessel development, as determined by positive staining for platelet endothelial cell adhesion molecule (PECAM-1), in the xenograft tumors (Figure 3D-F).

**Figure 3.**
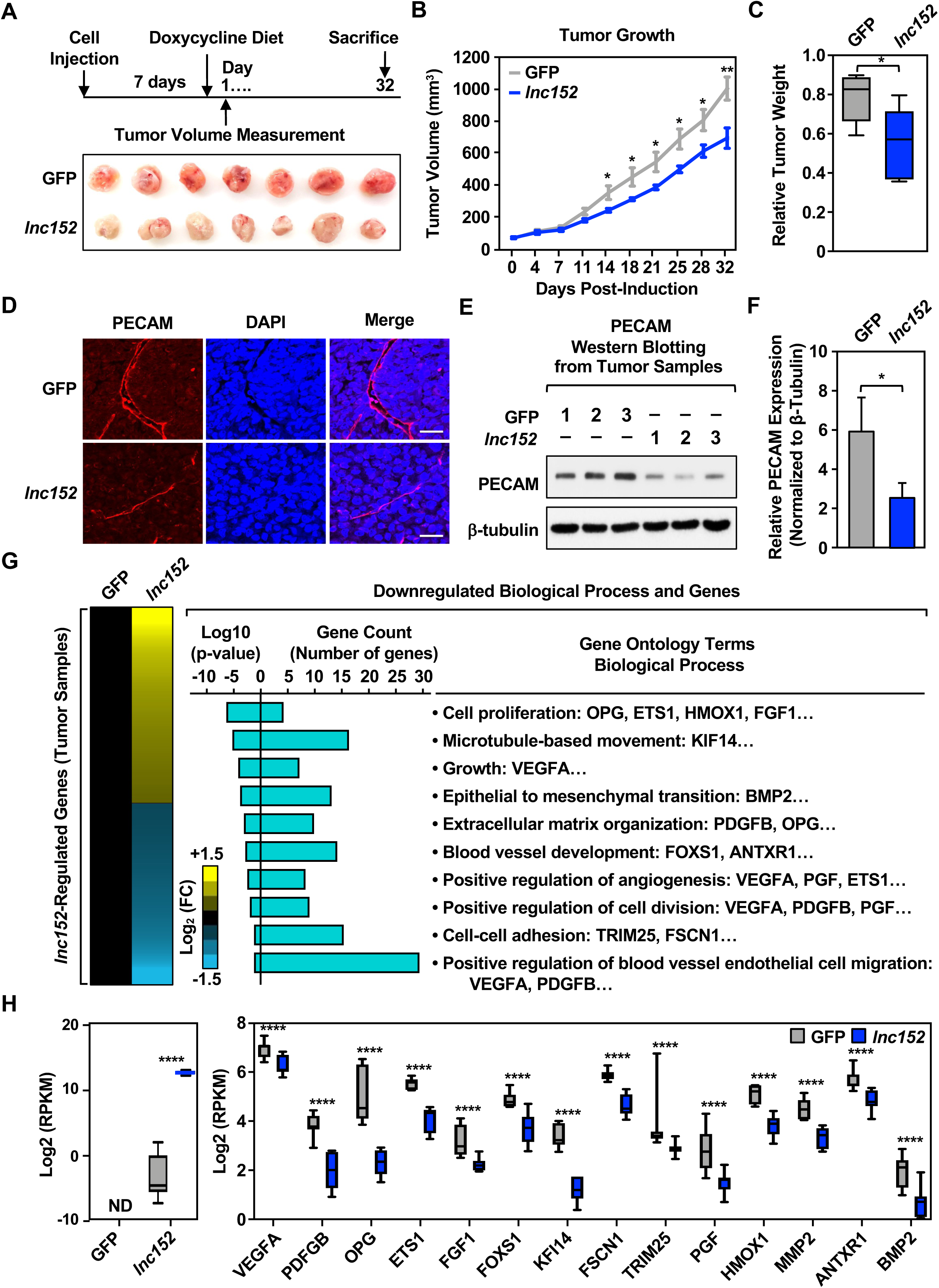
Ectopic expression of *lnc152* reduces tumor growth, inhibits angiogenesis, and alters gene expression in xenograft tumors in vivo. **(A)** Ectopic expression of *lnc152* inhibits the growth of MDA-MB-231 xenograft tumors. (*Top*) Schematic representation of the experimental timeline for the xenograft tumor model. (*Bottom*) Images of xenograft tumors formed from MDA-MB-231 cells expressing Dox-induced GFP or *lnc152*. **(B)** Growth curves for MDA-MB-231 xenograft tumors engineered for ectopic expression of GFP or *lnc152*. Each point represents the mean ± SEM, n = 7. Points marked with asterisks are statistically different from each other (two-tailed Student’s t-test, * p < 0.05, ** p < 0.01). **(C)** Box plots showing tumor weights at the end of the experiment. Points marked with asterisks are statistically different from each other (two-tailed Student’s t test, *p < 0.05). **(D-F)** Ectopic expression of *lnc152* inhibits blood vessel development in MDA-MB-231 xenograft tumors. (D) Representative immunohistochemical staining of PECAM from MDA-MB-231 xenograft tumor tissue from panel A. (E) Three tumor tissues from each group were analyzed by Western blotting for PECAM and β-tubulin. (F) Quantification of protein levels from the experiments shown in panel E. Bars marked with asterisks are statistically different from each other (two-tailed Student’s t test, * p < 0.05). **(G)** Gene expression and gene ontology analyses in MDA-MB-231 xenograft tumors. *(Left)* Heatmap showing the results of the RNA-seq assays from MDA-MB-231 tumors engineered for ectopic expression of GFP or *lnc15*. (*Right*) Gene ontology analysis showing terms and genes de-enriched upon ectopic *lnc152* expression in MDA-MB-231 xenograft tumors relative to ectopic expression of GFP. **(H)** Bar graphs showing the relative expression of a set of genes upon ectopic expression of *lnc152* from MDA-MB-231 xenograft tumor tissue (RNA-seq RPKM values, two-tailed Student’s t test, **** p < 0.001).

In RNA-seq assays, ectopic expression of *lnc152* altered the expression of hundreds of genes in the MDA-MB-231 xenograft tumors (Figure 3, G and H; Figure S3; Supplementary Tables S1 and S2). Interestingly, overexpression of *lnc152* caused downregulation of genes involved in biological processes such as cell proliferation, extracellular matrix organization, and blood vessel development (Figure 3G; Figure S4A). The global effects of *lnc152* on the downregulation of gene expression were reflected in the expression of individual genes controlling relevant biological processes (Figure 3H). These effects were confirmed by RT-qPCR from the same xenograft tumor tissues (Figure S3A), as well as in MDA-MB-231 cells (Figure S3B) and HCC1143 cells (Figure S3C). A set of genes were also upregulated upon ectopic expression of *lnc152* in the MDA-MB-231 xenografts (Figure S4B). Together, these results suggest that *lnc152* acts as a tumor suppressor in TNBC. Ectopic expression of *lnc152* reduces tumor growth, inhibits angiogenesis, and modulates the expression of genes that are involved in cell proliferation, blood vessel development, and positive regulation of angiogenesis in MDA-MB-231 xenograft tumors in vivo.

### *Lnc152* inhibits the metastasis of TNBC cells

The effects of *lnc152* on biological processes, such as cell proliferation, extracellular matrix organization, and blood vessel development, suggested a potential role for *lnc152* in metastasis. To test this possibility, we generated MDA-MB-231 cell lines stably expressing luciferase (Luc) with Dox-inducible ectopic expression of GFP or *lnc152* (Figure S5A) and used them in an intracardiac injection model of cancer metastasis (37) (Figure 4A). These Luc-expressing cell lines exhibited similar *lnc152*-dependent outcomes as the cell-based models shown above, such as cell proliferation (Figure S5B), effects on the expression of selected genes (Figure S5C), and tube formation in HUVECs on Matrigel (Figure S5, D and E).

**Figure 4.**
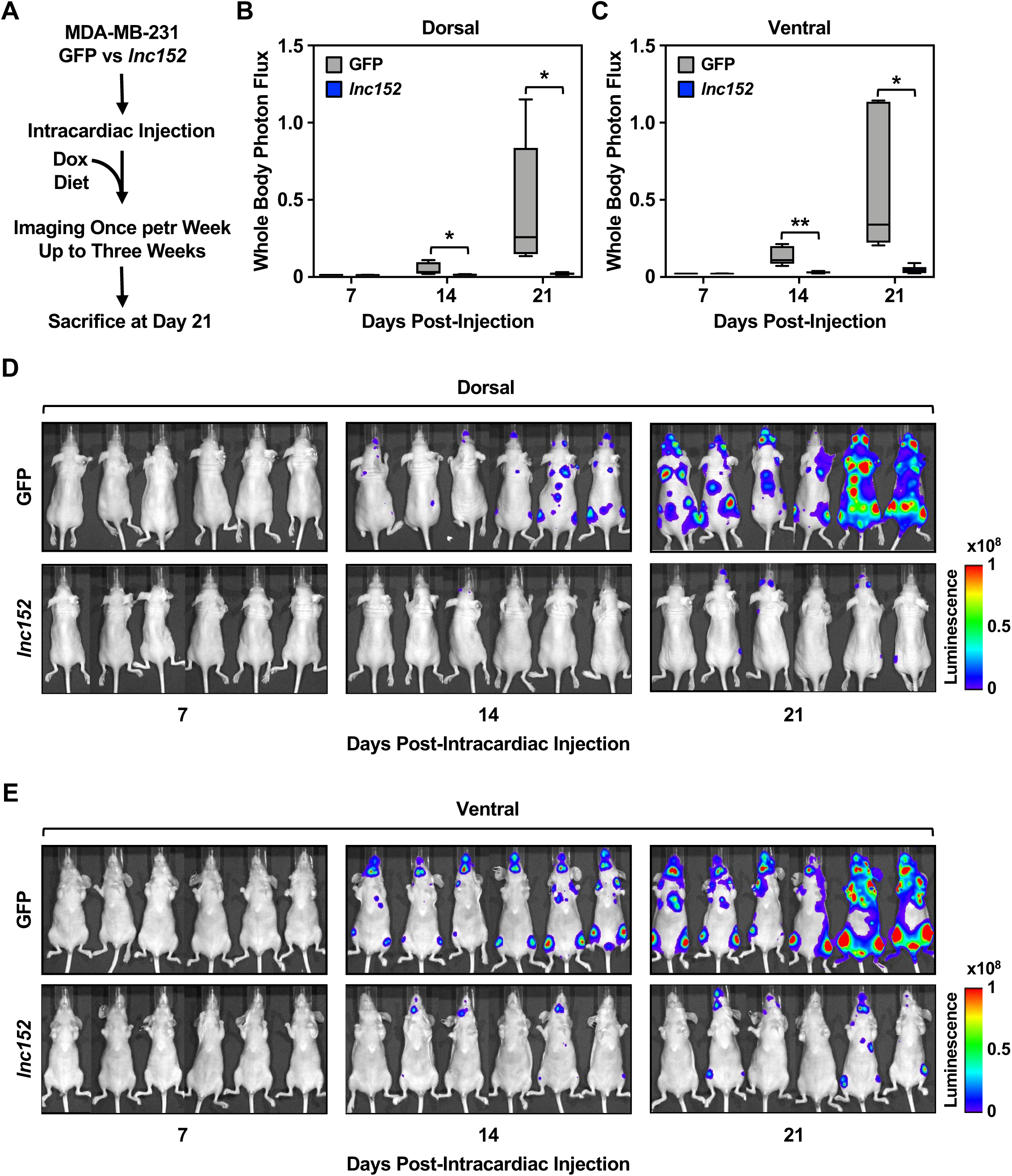
Ectopic expression of *lnc152* inhibits tumor metastasis in vivo. **(A)** Schematic representation of the experimental design for the intracardiac injection model of cancer metastasis in vivo. **(B-C)** Ectopic expression of *lnc152* reduces the metastatic burden in an MDA-MB-231 cell metastasis mouse model. Plot of bioluminescence signals from the dorsal (B) and ventral (C) views of mice at various times after intracardiac injection of tumor cells expressing GFP or *lnc152*. Each bar represents the mean ± SEM, n = 6. Bars marked with asterisks are statistically different from each other (two-tailed Student’s t-test, * p < 0.05, ** p < 0.01). **(D-E)** Bioluminescence tracing of the dorsal (D) and ventral (E) views from each group at various times after tumor cell intracardiac injection shows exponential increases in luciferase activity in mice expressing GFP compared to *lnc152*.

We used the Luc-expressing cell lines to examine the potential effects of *lnc152* on the metastasis of MDA-MB-231 cells. In this model, ectopic expression of *lnc152* inhibited metastasis of MDA-MB-231 cells to the brain, lung, and bones (Figure 4, B and C). Images of the bioluminescence signals from the dorsal (Figure 4D) and ventral (Figure 4E) views of the mice at various times after intracardiac injection of tumor cells shows a dramatic reduction in metastatic burden for cells expressing *lnc152*. In control experiments, dorsal bioluminescence signals from each group immediately after intracardiac injection show no significant differences in Luc activity in mice expressing *lnc152* compared to GFP (Figure S5, F-H). Together, these results demonstrate that expression of *lnc152* significantly modulates the metastatic behavior of TNBC cells.

### The RNA-binding tumor suppressor RBM47 interacts with *lnc152* and mediates similar functions

Interacting proteins are important effectors for lncRNA function. In turn, interactions with lncRNAs can modulate the stability and enzymatic activity of their interacting proteins. Thus, lncRNA-protein interactions can play a crucial role in fundamental cellular processes, including regulation of gene expression. To better understand the mechanism of action of *lnc152*, we sought to identify its interacting proteins in cells. In this regard, we used in vitro RNA-pull down assays coupled with LC-MS/MS analysis to determine the repertoire of *lnc152*-interacting proteins in MCF-7 breast cancer cells in which *lnc152* is naturally highly expressed. Proteomic analysis of the *lnc152* interactome identified a set of *lnc152*-interacting proteins (Figure 5, A-C; Figure S6, A and B; Supplementary Table S3), including RNA binding motif protein 47 (RBM47), which was one of the most highly enriched proteins in the *lnc152* interactome (Figure 5D). RBM47 is a known tumor suppressor and has been shown to inhibit breast cancer progression and metastasis (38,39). Furthermore, RBM47 alters the splicing and abundance of a subset of its target mRNAs (39). In vitro RNA-pull down assays confirmed that *lnc152* interacts directly with RBM47 (Figure 5, E-G).

**Figure 5.**
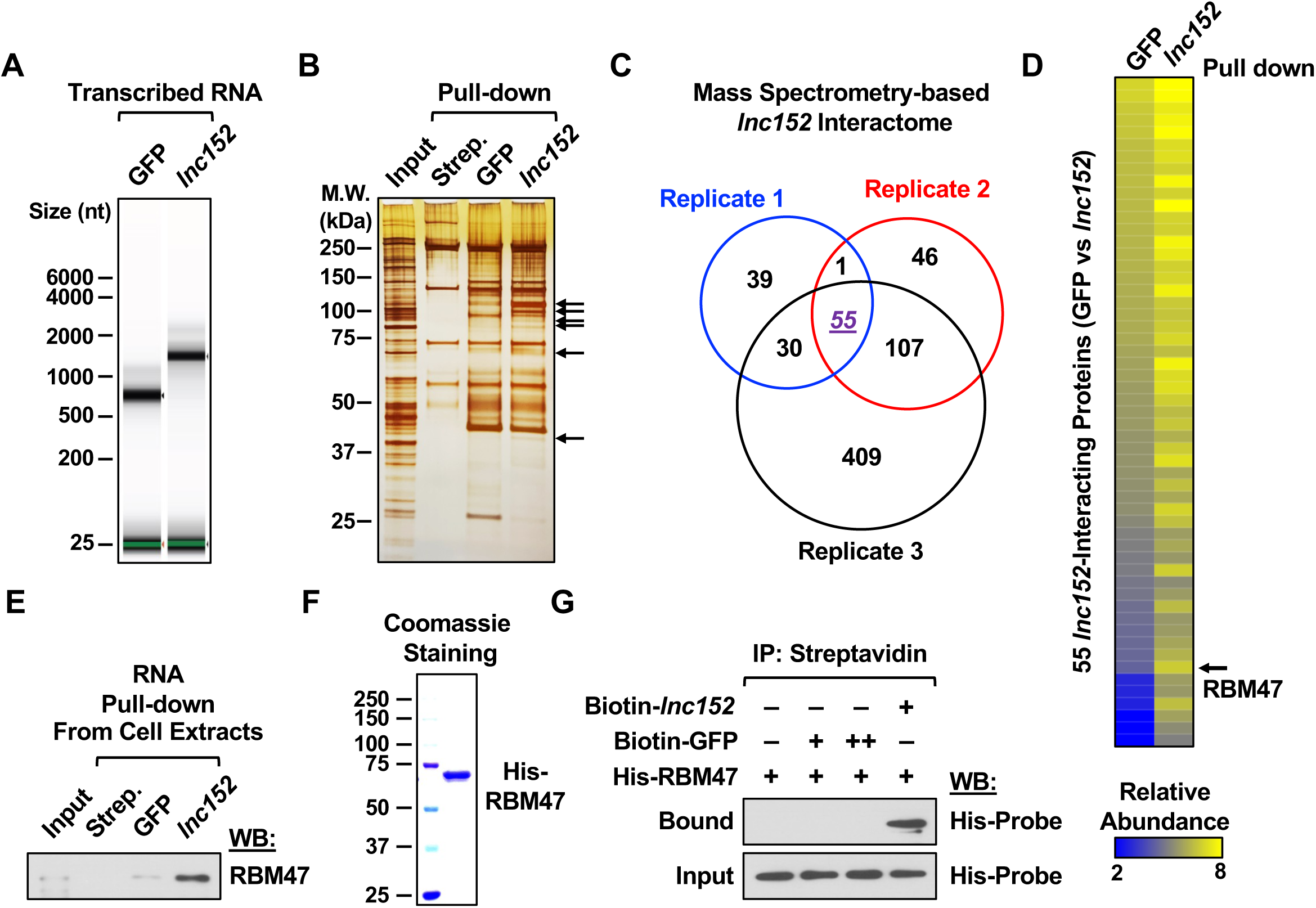
Proteomic analysis of the *lnc152* interactome reveals an association between *lnc152* and RBM47. **(A)** In vitro transcribed and biotinylated *GFP* or *lnc152* RNAs were visualized using an RNA BioAnalyzer. **(B)** Silver stained SDS-PAGE gel of proteins pulled down by *GFP* or *lnc152* RNAs from MCF-7 cell extracts. The arrows indicate significantly enriched proteins in the *lnc152* RNA pulldown compared to the *GFP* RNA pulldown. Streptavidin (Strep.) pulldown was used as a negative control. **(C)** A diagram depicting the *lnc152* interactome from three independent RNA pulldown assays. **(D)** Heatmap representing 55 *lnc152*-interacting proteins shared between three independent RNA pulldown assays (highlighted in purple in panel C). RBM47 is one of the most highly enriched proteins in the *lnc152* pulldown. **(E)** RNA pulldown assay showing a direct interaction between *lnc152* and RBM47. In vitro and biotinylated *GFP* or *lnc152* RNAs were incubated with whole cell extracts from MCF-7 cells, immunoprecipitated using streptavidin agarose beads, and then analyzed by Western blotting for RBM47. **(F)** Purified recombinant RBM47 protein was analyzed using SDS-PAGE with Coomassie blue staining. **(G)** RNA pulldown assay showing a direct interaction between *lnc152* and RBM47. In vitro transcribed and biotinylated *GFP* or *lnc152* RNAs were incubated with purified recombinant RBM47 protein, immunoprecipitated using streptavidin agarose beads, and then analyzed by Western blotting for RBM47.

To determine potential biological outcomes of *lnc152*-RBM47 interactions, we first determined if the expression of RBM47 correlates with the expression of *lnc152* across breast cancer subtypes or cell lines. We found that RBM47 is highly expressed in breast cancer tissues compared to normal breast tissues, based on data in TCGA (Figure 6A). RBM47 is upregulated in luminal and HER2-enriched tumors, but downregulated in basal-like tumors, from breast cancer patients (Figure 6B). Likewise, RBM47 is upregulated in luminal A, but downregulated in basal, breast cancer cell lines (Figure 6C).

**Figure 6.**
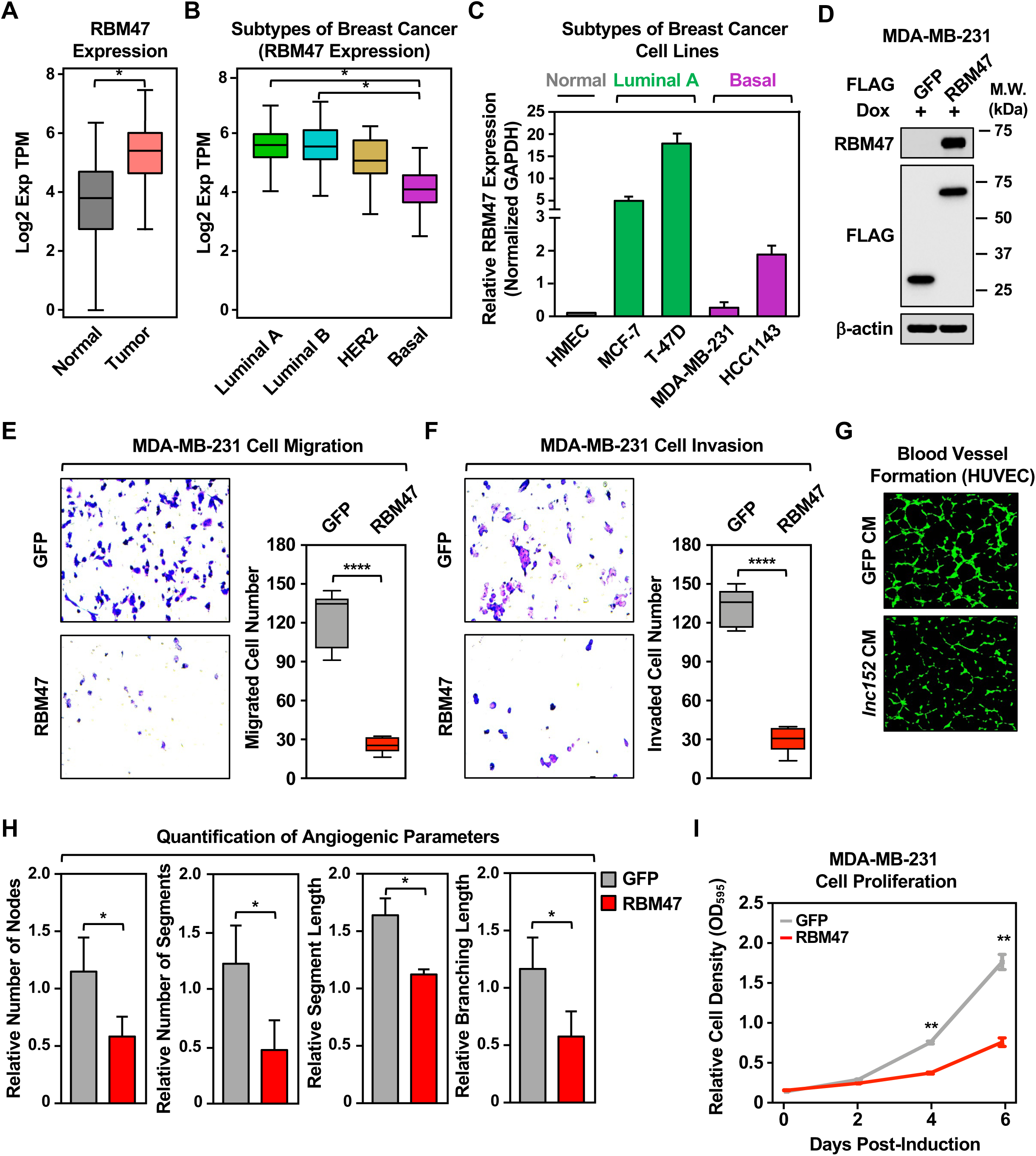
Ectopic expression of RBM47 inhibits aggressive cancer-related outcomes in MDA-MB-231 TNBC cells. **(A)** RNA-seq expression data for RBM47 in primary tumors (TCGA breast cancer samples, n = 1085) compared with normal breast tissues (TCGA normal and GTEx data, n = 291). Bars marked with asterisks are statistically different from each other (two-tailed Student’s t test, * p < 0.05). **(B)** Box plots comparing the expression of RBM47 in different intrinsic molecular subtypes of breast cancers [luminal A (n = 415), luminal B (n = 194), HER2-enriched (n = 66), and basal-like (n = 135)]. Bars marked with asterisks are statistically different from each other (two-tailed Student’s t test, * p < 0.05). **(C)** Bar graph showing the expression of RBM47 in breast cancer cell lines representing different molecular subtypes. **(D)** Generation of MDA-MB-231 cell lines for Dox-inducible ectopic expression of FLAG-GFP or FLAG-RBM47. Whole cell extracts were analyzed by Western blotting for RBM47, FLAG, and β-actin. Comparable protein expression of GFP and RBM47 levels are shown in the Western blots for FLAG. **(E)** (*Left*) Dox-induced ectopic expression of *lnc152* inhibits the migration of MDA-MB-231 cells compared to ectopic expression of GFP mRNA. (*Right*) Quantification of the results from the experiments shown in the left panel. Each bar represents the mean ± SEM, n = 6. Bars marked with asterisks are statistically different from each other (two-tailed Student’s t test, **** p < 0.0001). **(F)** (*Left*) Dox-induced ectopic expression of *lnc152* inhibits the invasion of MDA-MB-231 cells compared to ectopic expression of GFP mRNA. (*Right*) Quantification of the results from the experiments shown in the left panel. Each bar represents the mean ± SEM, n = 5. Bars marked with asterisks are statistically different from each other (two-tailed Student’s t test, **** p < 0.0001). **(G)** Dox-induced ectopic expression of RBM47 prevents human umbilical vein endothelial cell (HUVEC) tube formation on Matrigel. Morphological appearance of HUVECs grown on Matrigel with conditioned medium collected from MDA-MB-231 cells expressing Dox-induced GFP or RBM47, stained with Calcein-AM (green) and detected by fluorescence microscopy. **(H)** Quantification of the results from the experiments shown in panel G. Image J software with the Angiogenesis plugin was used to detect the number of nodes, number of segments, segment length, and branching length. Each bar represents the mean + SEM, n = 4. Bars marked with asterisks are statistically different from each other (two-tailed Student’s t test, * p < 0.05). **(I)** Dox-induced ectopic expression of RBM47 inhibits MDA-MB-231 cell proliferation more effectively than ectopic expression of GFP. Cells were treated daily with 1 μg/mL of Dox for six days. Each point represents the mean ± SEM, n = 3. Points marked with asterisks are statistically different from each other (two-tailed Student’s t-test, ** p < 0.01).

We also determined if ectopic expression of RBM47 might suppress the aggressive cancer features of TNBC lines, as seen with *lnc152* expression. Dox-induced ectopic expression of RBM47 (Figure 6D) inhibited the migration and invasion of MDA-MB-231 cells compared to ectopic expression of GFP (Figure 6, E and F). Moreover, Dox-induced ectopic expression of RBM47 prevented HUVEC tube formation on Matrigel using MDA-MB-231 conditioned medium (Figure 6, G and H). Finally, Dox-induced ectopic expression of RMB47 inhibited MDA-MB-231 cell proliferation compared to ectopic expression of GFP (Figure 6I). Together, these results illustrate similar expression patterns, roles, and functional outcomes for RBM47 and *lnc152*.

### *Lnc152* enhances RBM47 expression to drive cancer-related phenotypes

Both *lnc152* and RBM47 are downregulated in basal breast cancer cell lines, and ectopic expression of either *lnc152* or RBM47 suppressed the aggressive cancer phenotypes of TNBC cells, indicating that they are positively correlated in the regulation of TNBC. Since lncRNAs can promote gene expression and bind to proteins to regulate their stability, we tested the possibility that *lnc152* may affect *RBM47* mRNA expression or RBM47 protein stability. siRNA-mediated knockdown of *lnc152* decreased the levels RBM47 protein in MCF-7 cells (Figure 7A), whereas Dox-induced ectopic expression of *lnc152* increased the levels of RBM47 protein in both MDA-MB-231 cells (Figure 7B) and MDA-MB-231 xenograft tumor tissue (Figure 7C). Likewise, siRNA-mediated knockdown of *lnc152* decreased the steady-state levels of RBM47 mRNA in MCF-7 cells (Figure 7D), whereas Dox-induced ectopic expression of *lnc152* increased the levels of RBM47 mRNA in both MDA-MB-231 cells (Figure 7E) and MDA-MB-231 xenograft tumor tissue (Figure 7F). Furthermore, Dox-induced ectopic expression of *lnc152* in MDA-MB-231 xenograft tumor tissue increased the expression of RBM47-upregulated genes, such as *CASP7* and *HTATIP2*, which also act as tumor suppressors (Figure 7G; Figure S7, A and B) and decreased the expression levels of RBM47-downregulated genes, such as *SLC29A1, CDK2*, and *SUPT16H* (Figure 7G; Figure S7, A and B). Together, these results indicate that *lnc152* enhances RBM47 expression to drive cancer-related phenotypes.

**Figure 7.**
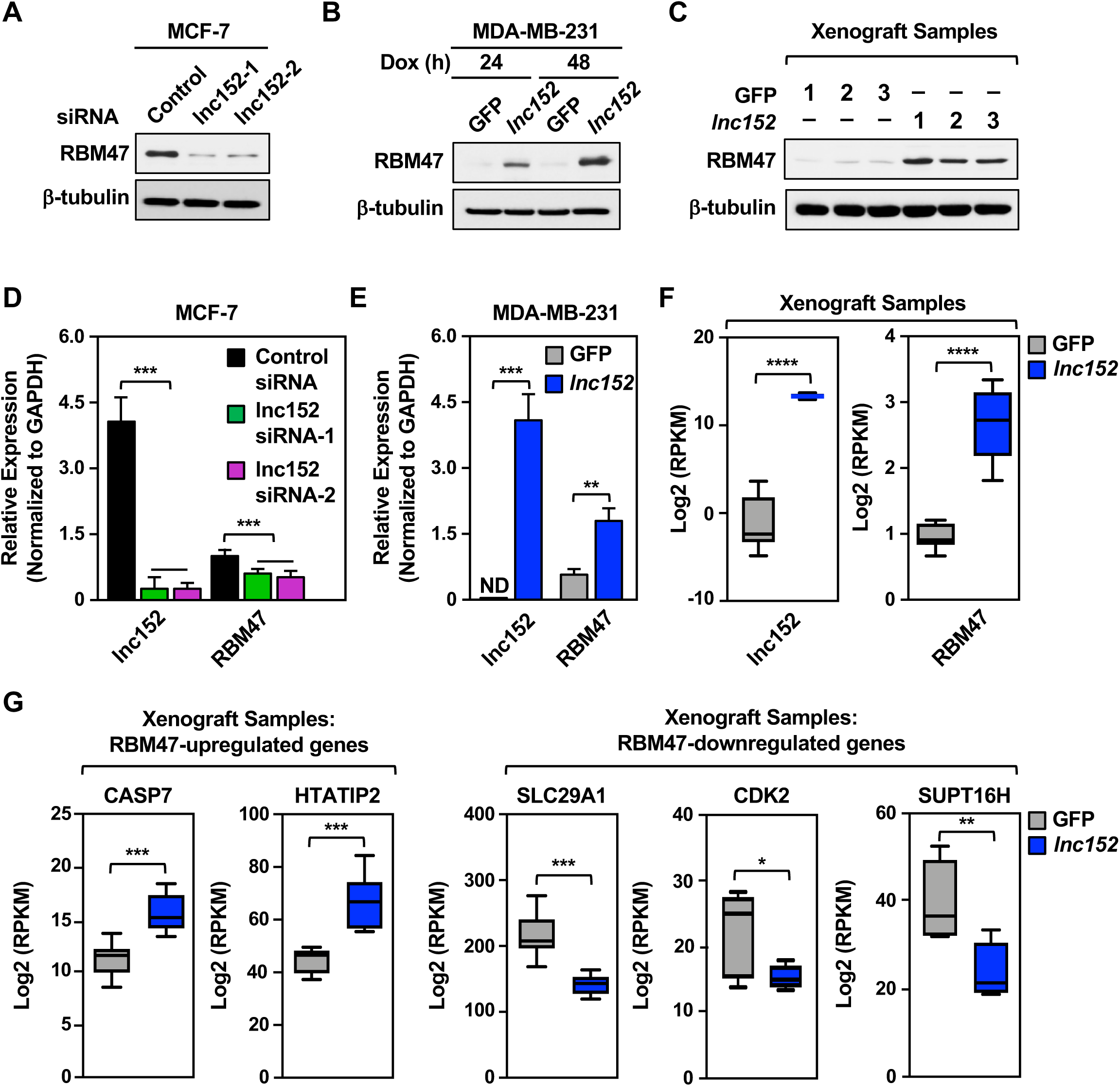
*Lnc152* stabilizes RBM47 mRNA and enhances RBM47 protein levels to promote RBM47-mediated gene expression in MDA-MB-231 TNBC cells. **(A)** siRNA-mediated knockdown of *lnc152* decreases the level of RBM47 in MCF-7 cells. Western blotting for RBM47 and β-tubulin is shown. **(B)** Dox-induced ectopic expression of *lnc152* increases the level of RBM47 in MDA-MB-231 cells. Western blotting for RBM47 and β-tubulin is shown. **(C)** Dox-induced ectopic expression of *lnc152* increases the level of RBM47 in MDA-MB-231 xenograft tumor tissue. Western blotting for RBM47 and β-tubulin is shown. **(D)** siRNA-mediated knockdown of *lnc152* decreases the steady-state levels of RBM47 mRNA in MCF-7 cells. *Lnc152* and RBM47 mRNA levels were determined by RT-qPCR and normalized to GAPDH mRNA. Each bar represents the mean + SEM, n = 3. Bars marked with asterisks are statistically different from each other (two-tailed Student’s t test, *** p < 0.001). **(E)** Dox-induced ectopic expression of *lnc152* increases the steady-state levels of RBM47 mRNA in MDA-MB-231 cells. *Lnc152* and RBM47 mRNA levels were determined by RT-qPCR and normalized to GAPDH mRNA. ND indicates that the expression level was not detectable. Each bar represents the mean + SEM, n = 3. Bars marked with asterisks are statistically different from each other (two-tailed Student’s t test, **p < 0.01, *** p < 0.001). **(F)** Bar graphs showing the relative expression of *lnc152* and RBM47 mRNA upon ectopic expression of *lnc152* in MDA-MB-231 xenograft tumor tissues, RNA-seq RPKM values (two-tailed Student’s t test, **** p < 0.0001). **(G)** Dox-induced ectopic expression of *lnc152* in MDA-MB-231 xenograft tumor tissue alters the expression levels of RBM47-regulated genes. Bar graphs showing the relative expression of RBM47-regulated genes, RNA-seq RPKM values (two-tailed Student’s t test, *p < 0.05, **p < 0.01, *** p < 0.001).

## Discussion

In the studies herein, we explored the cancer-related molecular functions of *lnc152* (a.k.a. *DRAIC*). Our results revealed some key features of the biology of *lnc152* in breast cancer. We found that *lnc152* is highly expressed in luminal breast cancer cells, where it acts to attenuate cell migration and invasion. Likewise, ectopic expression of *lnc152* inhibits cancer-related cellular phenotypes in TNBC cells, including angiogenesis and metastasis. Molecularly, *lnc152* interacts with RNA-binding tumor suppressor RBM47 and enhances RBM47 expression to mediate its inhibitory functions. Collectively, our results demonstrate that *lnc152* is an angiogenesis-inhibiting tumor suppressor that attenuates the aggressive cancer-related phenotypes found in TNBC.

### *lnc152/DRAIC* is emerging as an important regulator of cancer phenotypes

In a previous study, we annotated *lnc152* and characterized its functions in the control of cell cycle gene expression and proliferation in breast cancer cells (22). We observed effects of *lnc152* in proliferation, cell cycle progression, and regulation of the estrogen signaling pathway (22). A contemporaneous study identified *lnc152/DRAIC* as a tumor suppressor in prostate cancer, which is downregulated as the cells progress from an androgen-dependent to a castration-resistant state (23). Moreover, higher levels of *lnc152/DRAIC* in prostate cancers were found to be associated with longer disease-free survival (23). Subsequent studies have affirmed a key role for *lnc152/DRAIC* in the biology of an array of cancers, including breast, prostate, esophageal, and gastric cancers, as well as gliomas (24-30).

A wide range of molecular and cellular regulatory functions have been ascribed to *lnc152/DRAIC*, including gene regulation, inhibition of NF-κB activation, interference with NFRKB deubiquitylation, sponging miR-432-5p, inhibition of protein translation, and autophagy (24-30). These regulatory functions underlie the tumor suppressor functions of *lnc152/DRAIC*, as well as its role in regulating a range of cancer-related phenotypes, such as proliferation, migration, invasion, metastasis, and angiogenesis. Since the expression of *lnc152/DRAIC* varies dramatically in different cancer subtypes derived from the same tissue, it may have value as a biomarker or prognostic indicator.

### Functional interactions between lnc152 and RBM47

One mechanism by which lncRNAs exert their regulatory effects is through specific interacting proteins. Using in vitro RNA-pull down assays coupled with LC-MS/MS, we identified the RNA-binding tumor suppressor RBM47 as a *lnc152*-interacting protein. Previous studies have shown that RBM47 suppresses cancer phenotypes, including proliferation, migration, invasion, and metastasis, in breast cancer and non-small-cell lung cancer (NSCLC) by regulating the splicing and stabilization of its target mRNAs (38,39). For example, RBM47 binds to the 3′-UTR of *DKK1* mRNA, which encodes DKK1 (Dickkopf Wnt signaling pathway inhibitor 1), to protect it from destabilizing factors, thus increasing DKK1 secretion in TNBC (39). RBM47 also binds the *AXIN1* mRNA, which encodes a pro-oncogenic β-catenin signaling pathway regulator, thereby stabilizing it and enhancing suppression of Wnt/β-catenin signaling in NSCLC (38). We speculate that RBM47 uses the same features that allow it to bind to mRNAs to bind to *lnc152*.

Although we have not yet elucidated the precise molecular underpinnings of the functional interactions between *lnc152* and RBM47, we have shown that the effects of *lnc152* in TNBC cells are mediated, in part, by regulating the expression of RBM47. *Lnc152* elevates the expression of RBM47 mRNA and protein, which impacts a wide range of cancer-related outcomes. Interestingly, *lnc152* and RBM47 are both highly expressed in the luminal subtype of breast cancer, and significantly downregulated in the triple-negative subtype, suggesting that *lnc152* and RBM47 levels may serve as biomarkers to predict aggressive features of breast cancer.

### *lnc152* inhibits angiogenesis

Angiogenesis is an essential component of the metastatic pathway. The process begins with invasion and intravasation into the surrounding lymphatics and blood vessels, and culminates with colonization of the disseminated tumor cells in a distal organ, angiogenesis, and growth (1,2). Our transcriptome analysis of TNBC cells xenografts indicated that *lnc152* downregulates genes controlling angiogenesis. Moreover, we found that ectopic expression of *lnc152* inhibits angiogenic phenotypes in HUVECs by TNBC cells whose proliferation is inhibited by *lnc152* expression (e.g., MDA-MB-231 cells) likely through the production of a soluble factor. Such an inhibitory effect of *lnc152* on angiogenesis could explain the potent inhibition of TNBC cell metastasis that we observed in mice. In this regard, new blood vessels promote tumor metastasis by providing the route of exit for tumor cells from the primary tumor and colony formation at secondary sites (3,4). Moreover, new blood vessels within the tumor are needed to provide sufficient oxygen and nutrients to support tumor growth (3,4).

Collectively, our results identify *lncRNA152* as an angiogenesis-inhibiting tumor suppressor that attenuates the aggressive cancer-related phenotypes found in TNBC by upregulating the expression of tumor the suppressor RBM47. As such, *lncRNA152* may serve as a biomarker to track aggressiveness of breast cancer, as well as therapeutic target for treating TNBC.

## Materials and Methods

Additional details about the materials and methods can be found in the Supplementary Materials and Methods under the same headings listed here.

### Cell Culture and Treatments

MCF-7, MDA-MB-231, T-47D, HCC1143, 293T, primary human mammary epithelial cells (HMEC), and human umbilical vein endothelial cells (HUVEC) cells were purchased from the American Type Cell Culture (ATCC). Luciferase-expressing MDA-MB-231 cells (40) were kindly provided by Dr. Srinivas Malladi. A detailed description of the culture conditions is provided in the Supplementary Materials and Methods. Fresh cell stocks were regularly replenished from the original stocks, verified for cell type identity using the GenePrint 24 system (Promega, B1870), and confirmed as mycoplasma-free every three months using a commercial testing kit. For induction with doxycycline (Dox; Sigma-Aldrich, D9891), the cells were treated with 0.5-1 μg/mL of Dox for the indicated times before collection.

### Molecular Cloning to Generate Expression Vectors

cDNA pools were prepared by extraction of total RNA from MCF-7 cells as described previously (41) to amplify *lnc152* and RBM47 cDNAs, which were then cloned into multiple vectors such as pcDNA3, pET19b, and pINDUCER20.

### Purification of RBM47 Protein Expressed in *E. coli*

His-tagged human RBM47 was expressed in *E. coli* Rosetta (DE3) cells using the pET19b-based bacterial expression vector. The transformed bacteria were grown in LB containing ampicillin and chloramphenicol at 37 °C until the OD595 reached ∼0.7. Recombinant protein expression was induced by the addition of 0.5 mM IPTG for 14 h at 20 °C. The cells were collected by centrifugation, and the cell pellets were flash-frozen in liquid N_2_. RBM47 was purified from the cells as described previously (41). The purified proteins were quantified using a Bradford protein assay (Bio-Rad), aliquoted, flash-frozen in liquid N_2_, and stored at −80°C.

### Generation of Knockdown, Knockout, and Ectopic Expression Cell Lines

#### siRNA-mediated knockdown in MCF-7 cells

siRNA oligos targeting human *lnc152* [described previously (22)] were transfected into MCF-7 cells at a final concentration of 10 nM using Lipofectamine RNAiMAX reagent (Invitrogen, 13778150) according to the manufacturer’s instructions.

#### Dox-inducible ectopic expression in MDA-MB-231 and HCC1143 cells

Lentiviruses were generated by transfection of pINDUCER20-based vectors (Addgene, plasmid no. 44012) for Dox-inducible expression of *lnc152*, GFP, or RBM47, along with pCMV-VSV-G, pCMV-GAG-Pol-Rev, and pAdVAntage into 293T cells using Lipofectamine 3000 according to the manufacturer’s protocol. Stably transduced cells were isolated under drug selection with 1 μg/mL Geneticin (Life Technologies, 11811031), and were used for Dox-induced ectopic expression of *lnc152*, GFP, and RBM47 to perform a variety of experiments described herein.

### Cell Proliferation Assays

Cells expressing *lnc152*, GFP, or RBM47 were plated and grown with daily changes of Dox-containing medium (0.5 μg/mL) for the indicated times before collection. Cell proliferation assays were performed on fixed cells using a crystal violet staining assay. After washing to remove unincorporated stain, the crystal violet was extracted using 10% glacial acetic acid and the absorbance was read at 595 nm.

### Preparation of Nuclear Extracts and Western Blotting

Whole cell lysates from cell lines and xenografts were prepared as described previously (41). Western bolting from cell and tumor tissue extracts were performed as described previously (41).

### RNA Isolation and Reverse Transcription Quantitative PCR (RT-qPCR)

The cells were collected at the indicated time points and cDNA pools were prepared by extraction of total RNA from cell lines using Trizol (Sigma-Aldrich, T9424), followed by reverse transcription using MMLV reverse transcriptase (Promega, M150B) with random hexamer or oligo(dT) primers (Sigma-Aldrich). The cDNA samples were subjected to qPCR as described previously (41) using gene-specific primers.

### RNA-sequencing

#### Generation of RNA-seq libraries

Total RNA from seven xenograft tumor tissues of each group was extracted using the RNeasy Plus Kit (Qiagen, 74134) and used to generate strand-specific RNA-seq libraries as described previously (42). The RNA-seq libraries were subjected to QC analyses (i.e., number of PCR cycles required to amplify each library, the final library yield, and the size distribution of the final library DNA fragments) and sequenced using an Illumina HiSeq 2500 and NextSeq 500.

#### Analysis of RNA-seq data sets

The raw data were subjected to QC analyses using the FastQC (43). The raw reads were aligned to the human reference genome (hg19/GRCh37) using default parameters in Tophat (v2.0.12) (44). Uniquely mappable reads were converted into bigWig files using BEDTools (45) for visualization in the Integrative Genomics Viewer (46). Transcriptome assembly was performed using cufflinks v.2.2. (47) with default parameters. The transcripts were merged into distinct, non-overlapping sets using cuffmerge, followed by cuffdiff to call the differentially regulated transcripts (47).

#### Gene set enrichment analysis

Gene ontology (GO) analyses were performed using the DAVID (Database for Annotation, Visualization, and Integrated Discovery) tool (48).

### Mining of Public Databases

The expression of *lnc152* and RBM47 in normal and different subtypes of breast cancer tissues was determined using GEPIA based on the RPKM values in the TCGA dataset (49).

### Analysis of ChIP-seq Data

FoxA1 ChIP-seq libraries from MCF-7 cells (NCBI GEO accession number GSE59530) were generated and analyzed as described previously (50).

### In Vitro Transcription to Generate GFP mRNA and *lnc152*

The in vitro transcribed GFP mRNA and *lnc152* were prepared as described previously (41). To generate biotin-labeled RNAs for pull down assays, an NTP labeling mixture containing biotin-UTP (10 mM each ATP, CTP, and GTP, 6.5 mM UTP, 3.5 mM biotin-UTP; Sigma, 11685597910) was used for in vitro transcription.

### In Vitro RNA Pull Down Assays Combined with LC-MS/MS Analysis

#### In vitro RNA pull down

Whole cell extract from MCF-7 cells was incubated with 5 nM of folded RNA at room temperature for 1 hour, followed by incubation with equilibrated Streptavidin-agarose beads (ThermoFisher Scientific, 20377) at room temperature for 30 minutes. The beads containing the protein-RNA complex were washed four times in Binding Buffer containing an additional 200 mM of NaCl. The reactions were stopped by the addition of 4x SDS-PAGE Loading Buffer, followed by heating to 100°C for 10 minutes.

#### LC-MS/MS analysis

Gel purified samples were injected onto an Orbitrap Fusion Lumos mass spectrometer (Thermo Electron) coupled to an Ultimate 3000 RSLC-Nano liquid chromatography system (Dionex). A detailed description of sample processing is provided in the Supplementary Materials and Methods.

### Cell Migration and Invasion Assays

Boyden chamber assays were used to determine the migration and invasive capacity of cells. Cells that migrated or invaded into the lower side of the membrane were fixed and stained with 0.5% crystal violet in 20% methanol solution for 15 minutes, washed with water, and air-dried.

### In Vitro Endothelial Tube Formation Assay

#### Preparation of conditioned medium (CM)

The serum-free conditioned medium in the presence of 1 μg/mL of Dox was collected and concentrated using Centricon 10 (Millipore) centrifuge filter unit, aliquoted, flash-frozen in liquid N_2_, and stored at −80 °C.

#### Endothelial tube formation assay

HUVECs were resuspended in conditioned medium from MDA-MB-231 cells expressing GFP mRNA or *lnc152* and were seeded onto Matrigel Basement Membrane Matrix coated 24-well plates and incubated at 37°C for 4-12 hours. Capillary-like tubes were then stained with cell-permeable dye Calcein AM (ThermoFisher, C1430) and imaged by fluorescence microscopy.

#### Quantification of tube networks

The total number of nodes, number of segments, total segment length, and branching were analyzed by the Angiogenesis Analyzer plugin in ImageJ.

### Indirect Immunofluorescence

Xenograft tissues were processed for paraffin sectioning using standard protocols. After staining, the immunofluorescent signals were imaged using a Zeiss LSM880 confocal microscope.

### Experiments with Mice

All animal experiments were performed in compliance with the Institutional Animal Care and Use Committee (IACUC) at the UT Southwestern Medical Center. Female athymic nude (*Foxn1*^*nu*^; Envigo #069) mice at 6-8 weeks of age were used.

#### Xenograft experiment

Breast cancer xenografts using MDA-MB-231 cells engineered for Dox-inducible expression of GFP mRNA or *lnc152* were performed as described previously (41).

#### Metastasis experiments

MDA-MB-231 cells harboring a luciferase reporter were engineered for Dox-inducible expression of GFP mRNA or *lnc152* and injected intracardially into the arterial circulation of 5 weeks old female athymic nude (*Foxn1*^*nu*^) mice (Envigo #069) to mimic hematogenous dissemination (37). Successful injection was confirmed by the detection of luciferase signal in the whole mouse.

#### Whole-body bioluminescent imaging

Mice were monitored weekly for metastatic outgrowth by bioluminescence imaging. D-luciferin (150 mg/kg) was injected retro-orbitally, and the mice were imaged using IVIS Spectrum with Living Image 4.4 software (PerkinElmer).

### Data and Code Availability

#### Genomic datasets

The new RNA-seq data generated for this study can be accessed from the NCBI’s Gene Expression Omnibus (GEO) repository (http://www.ncbi.nlm.nih.gov/geo/) using the superseries accession number GSE193634.

#### Custom scripts

Custom R scripts for genomic data analyses are available from the Lead Contact on request.

#### Mass spectrometry datasets

The new mass spectrometry datasets generated for this study can be accessed from MassIVE and also are available as supplemental data provided with this manuscript.

## Supporting information

Supplemental materials (figures, tables, methods, references)

Table S1 - Lnc152-downregulated genes in MDA-MB-231 xenograft tumors

Table S2 - Lnc152-upregulated genes in MDA-MB-231 xenograft tumors

Table S3 - Lnc152-interacting proteins

## Supplementary Information

This paper includes supplementary tables, figures, materials and methods, and references.

## Acknowledgements

We thank members of the Kraus lab for intellectual input and critical comments on this manuscript and Anusha Nagari for assistance with analysis of sequencing data. We acknowledge and thank the following UT Southwestern core facilities: Live Cell Imaging Core for microscopy (Dr. Katherine Luby-Phelps; NIH S10 OD021684), Next Generation Sequencing Core for deep sequencing (Dr. Ralf Kittler), Proteomics Core Facility for mass spectrometry (Dr. Andrew Lemoff), Histo Pathology Core for histology (Dr. Bret Evers), and the Small Animal Imaging Shared Resource, which is supported in part by the Harold C. Simmons Cancer Center through an NCI Cancer Center Support Grant (P30 CA142543).

## Funding

This work was supported by grants from the Cancer Prevention and Research Institute of Texas (RP130607, RP160318, RP190235) and funds from the Cecil H. and Ida Green Center for Reproductive Biology Sciences Endowment to W.L.K.; a postdoctoral fellowship from the U.S. Department of Defense Breast Cancer Research Program (BCRP) to D.S.K. (BC134066); and a Komen for the Cure Foundation postdoctoral fellowship to S.S.G. (PDF230441).

## Author Contributions

D.S.K., S.S.G. and W.L.K. conceived this project, designed the experiments, and oversaw their execution. D.S.K. performed all of the biochemical and cell-based experiments. D.S.K. and S.S.G. performed the mouse xenograft experiments. D.S.K. prepared the RNA-seq libraries from tumor tissues. S.S.G. performed all of the FoxA1-related experiments. R.S. and T.N. analyzed the RNA-seq and mass spectrometry data. D.S.K., K.K., and S.M. performed the mouse metastasis experiments. T.H. analyzed RBM47-regulated genes from a genomic data set. D.S.K and C.V.C. prepared the initial drafts of the figures and text, which were edited and finalized by W.L.K. W.L.K. secured funding to support this project and provided intellectual support for all aspects of the work.

## Competing Interests

The authors have no competing interests to declare for this work.

## Notes

**<u>Funding</u>**: This work was supported by grants from the Cancer Prevention and Research Institute of Texas (RP130607, RP160318, RP190235) and funds from the Cecil H. and Ida Green Center for Reproductive Biology Sciences Endowment to W.L.K.; a postdoctoral fellowship from the U.S. Department of Defense Breast Cancer Research Program (BCRP) to D.S.K. (BC134066); and a Komen for the Cure Foundation postdoctoral fellowship to S.S.G. (PDF230441).

### Competing Interest Statement

The authors have declared no competing interest.

