## Supplemental materials (figures, tables, methods, references) for "Functional Characterization of *lncRNA152* as an Angiogenesis-Inhibiting Tumor Suppressor in Triple-Negative Breast Cancers"

**Kim *et al.* (2021)**

This document contains the following: Page

#### 1) SUPPLEMENTARY TABLES AND LEGENDS

**Supplementary Table S1. Lnc152-downregulated genes in MDA-MB-231 xenograft tumors.**

Total RNA from seven xenograft tumor tissues of each group was extracted and used to generate strand-specific RNA-seq libraries. The significantly ($q < 0.001$) downregulated genes were determined by comparing the ectopic expression of lnc152 samples to ectopic expression of GFP samples. The "Column Heading" key provides annotation information describing the identified gene sets. The "Tab" key provides details about the worksheets contained within this spreadsheet.

**Supplementary Table S2. Lnc152-upregulated genes in MDA-MB-231 xenograft tumors.**

Total RNA from seven xenograft tumor tissues of each group was extracted and used to generate strand-specific RNA-seq libraries. The significantly ($q < 0.001$) upregulated genes were determined by comparing the ectopic expression of lnc152 samples to ectopic expression of GFP samples. The "Column Heading" key provides annotation information describing the identified gene sets. The "Tab" key provides details about the worksheets contained within this spreadsheet.

**Supplementary Table S3. Lnc152-interacting proteins.**

GFP mRNA or *lnc152* RNA were transcribed and labeled with biotin in vitro and incubated with whole cell extract from MCF-7 cells. *Lnc152*-interacting proteins were identified by enrichment in the pulldown with *lnc152* RNA versus GFP mRNA ($Log_2$ FC). The "Column Heading" key provides annotation information describing the lnc152-interacting proteins. The "Tab" key provides details about the worksheets contained within this spreadsheet.

#### 2) SUPPLEMENTARY FIGURES AND LEGENDS

*[Figure S1 is on the next page]*

**Figure S1. FoxA1 regulates the expression of *lnc152*.**
**(A)** Overview of the computational analysis pipeline used for the identification of *lnc152* in MCF-7 cells.
**(B)** Genome browser tracks of RNA-seq and ChIP-seq data showing FoxA1 peaks within the promoter region of *lnc152*.
**(C)** Cumulative distribution curves showing the correlation in gene expression profiles of FoxA1 and *lnc152* in the TCGA breast cancer datasets.
**(D)** RNA-seq expression data from TCGA breast cancer datasets showing a strong positive correlation between FoxA1 expression and *lnc152* expression. RSEM = RNA-Seq by Expectation Maximization.
**(E)** RNA-seq expression data from GTEx normal breast datasets showing a moderate positive correlation between FoxA1 expression and *lnc152* expression. RPKM = Reads Per Kilobase Million.
**(F)** RNA-seq expression data from CCLE datasets showing the correlation between key transcription factors (FoxA1, GAT3 or ESR1) and l*nc152*. RMA = Robust Multiarray Averaging.
**(G)** Bar graphs showing the expression of FoxA1 and *lnc152* in breast cancer cell lines representing different intrinsic molecular subtypes as quantified by RT-qPCR. ND indicates expression level was not detectable.
**(H)** siRNA-mediated knockdown of FoxA1 decreases the expression level of *lnc152* in MCF-7 and T-47D luminal breast cancer cell lines.

### Figure S1.

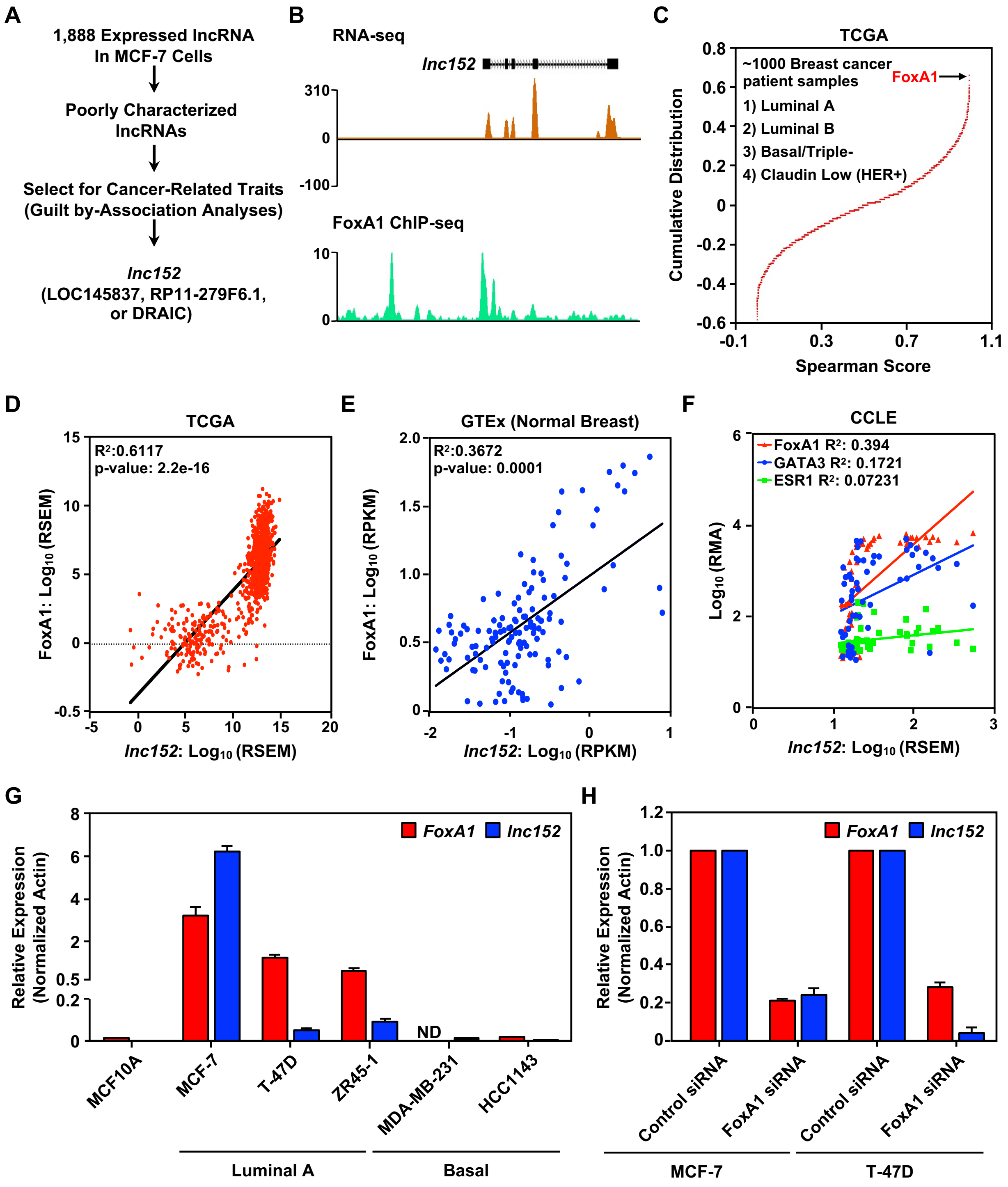
A
1,888 Expressed lncRNA In MCF-7 Cells
Poorly Characterized lncRNAs
Select for Cancer-Related Traits (Guilt by-Association Analyses)
lnc152 (LOC145837, RP11-279F6.1, or DRAIC)
B
RNA-seq
lnc152
FoxA1 ChIP-seq
C
TCGA
~1000 Breast cancer patient samples
1) Luminal A
2) Luminal B
3) Basal/Triple-
4) Claudin Low (HER+)
FoxA1
Cumulative Distribution
Spearman Score
D
TCGA
R²:0.6117
p-value: 2.2e-16
FoxA1: Log10 (RSEM)
lnc152: Log10 (RSEM)
E
GTEx (Normal Breast)
R²:0.3672
p-value: 0.0001
FoxA1: Log10 (RPKM)
lnc152: Log10 (RPKM)
F
CCLE
FoxA1 R²: 0.394
GATA3 R²: 0.1721
ESR1 R²: 0.07231
Log10 (RMA)
lnc152: Log10 (RSEM)
G
FoxA1
lnc152
Relative Expression (Normalized Actin)
ND
MCF10A
MCF-7
T-47D
ZR45-1
MDA-MB-231
HCC1143
Luminal A
Basal
H
FoxA1
lnc152
Relative Expression (Normalized Actin)
Control siRNA
FoxA1 siRNA
Control siRNA
FoxA1 siRNA
MCF-7
T-47D

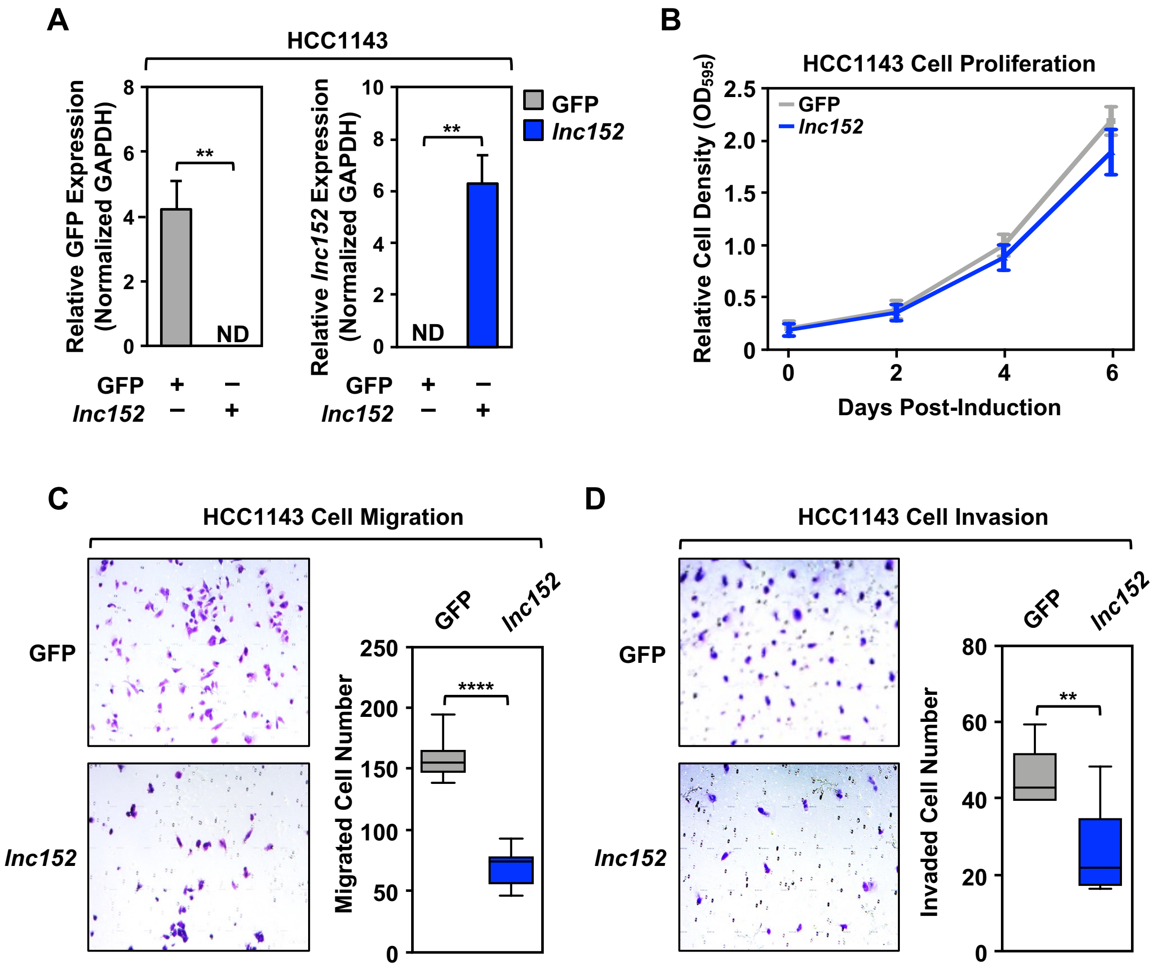

**Figure S2. Ectopic expression of *lnc152* inhibits aggressive cancer-related outcomes in HCC1143 TNBC cells.**

**(A)** Generation of HCC1143 cell lines for Dox-inducible ectopic expression of GFP or *lnc152*. GFP mRNA and *lnc152* levels were determined by RT-qPCR and normalized to GAPDH mRNA. ND indicates expression level was not detectable. Each bar represents the mean + SEM, n = 3. Bars marked with asterisks are statistically different from each other (two-tailed Student's t test, ** $p < 0.01$).

**(B)** Dox-induced ectopic expression of *lnc152* had no effect on HCC1143 cell proliferation. Cells were treated daily with 0.5 µg/mL of Dox for six days. Each point represents the mean ± SEM, n = 3.

**(C)** (*Left*) Dox-induced ectopic expression of *lnc152* inhibits the migration of HCC1143 cells compared to expression of GFP mRNA. (*Right*) Quantification of the results from the experiments shown in the left panel. Each bar represents the mean ± SEM, n = 7. Bars marked with asterisks are statistically different from each other (two-tailed Student's t test, **** $p < 0.0001$).

**(D)** (*Left*) Dox-induced ectopic expression of *lnc152* inhibits the invasion of HCC1143 cells compared to expression of GFP mRNA. (*Right*) Quantification of the results from the experiments shown in the left panel. Each bar represents the mean ± SEM, n = 6. Bars marked with asterisks are statistically different from each other (two-tailed Student's t test, ** $p < 0.01$).

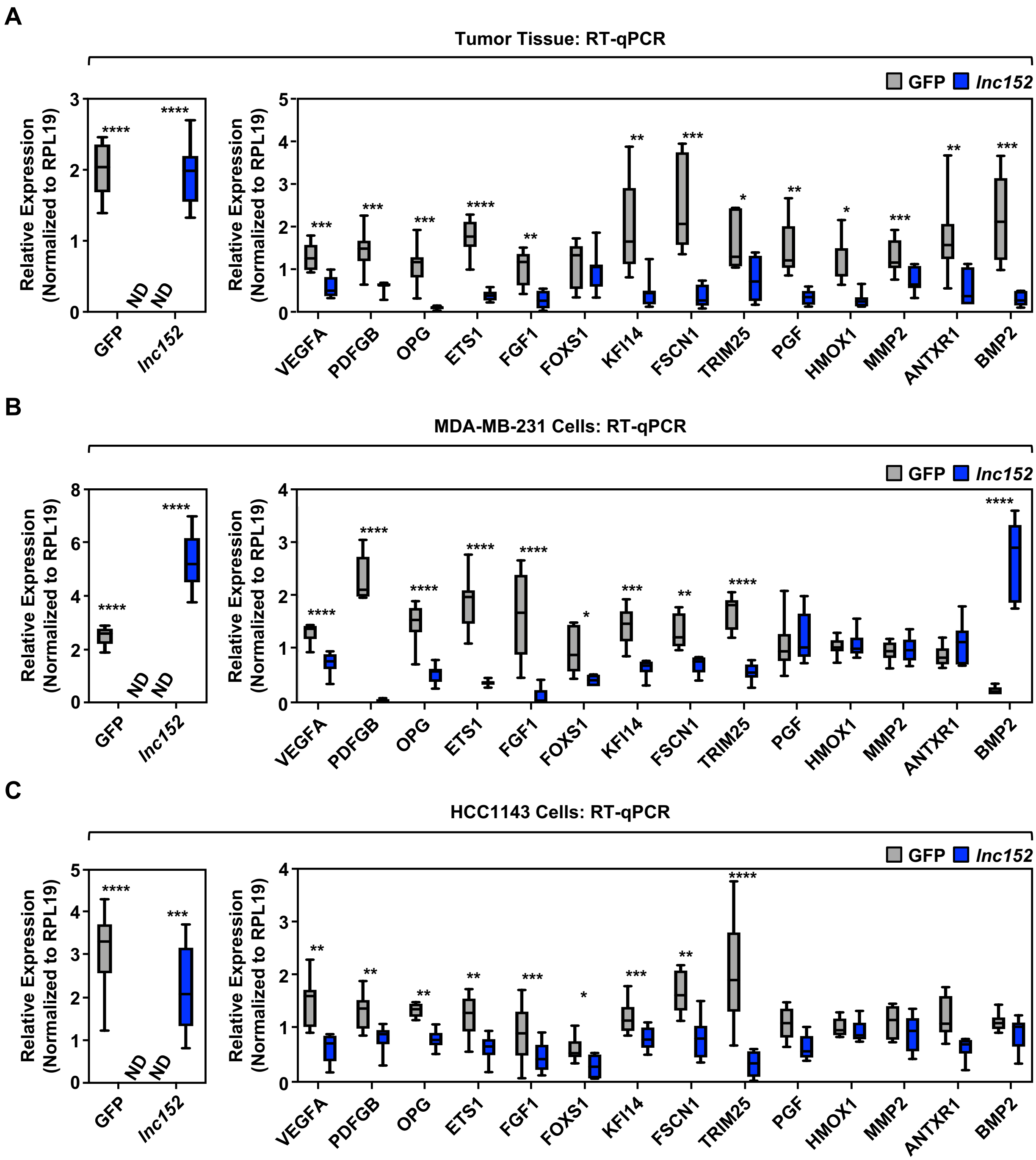

**Figure S3. Ectopic expression of *lnc152* alters cancer-related gene expression.**
**(A-C)** Bar graphs showing the expression levels of selected subsets of *lnc152*-regulated genes measured upon ectopic expression of *lnc152*, as assessed by RT-qPCR from xenograft tumor tissues (A), MDA-MB-231 cells (B), and HCC1143 cells (C), all normalized to RPL19 mRNA. ND indicates that expression was not detectable. Bars marked with asterisks are statistically different from each other (two-tailed Student's t test, * $p < 0.05$, ** $p < 0.01$, *** $p < 0.001$, **** $p < 0.0001$).

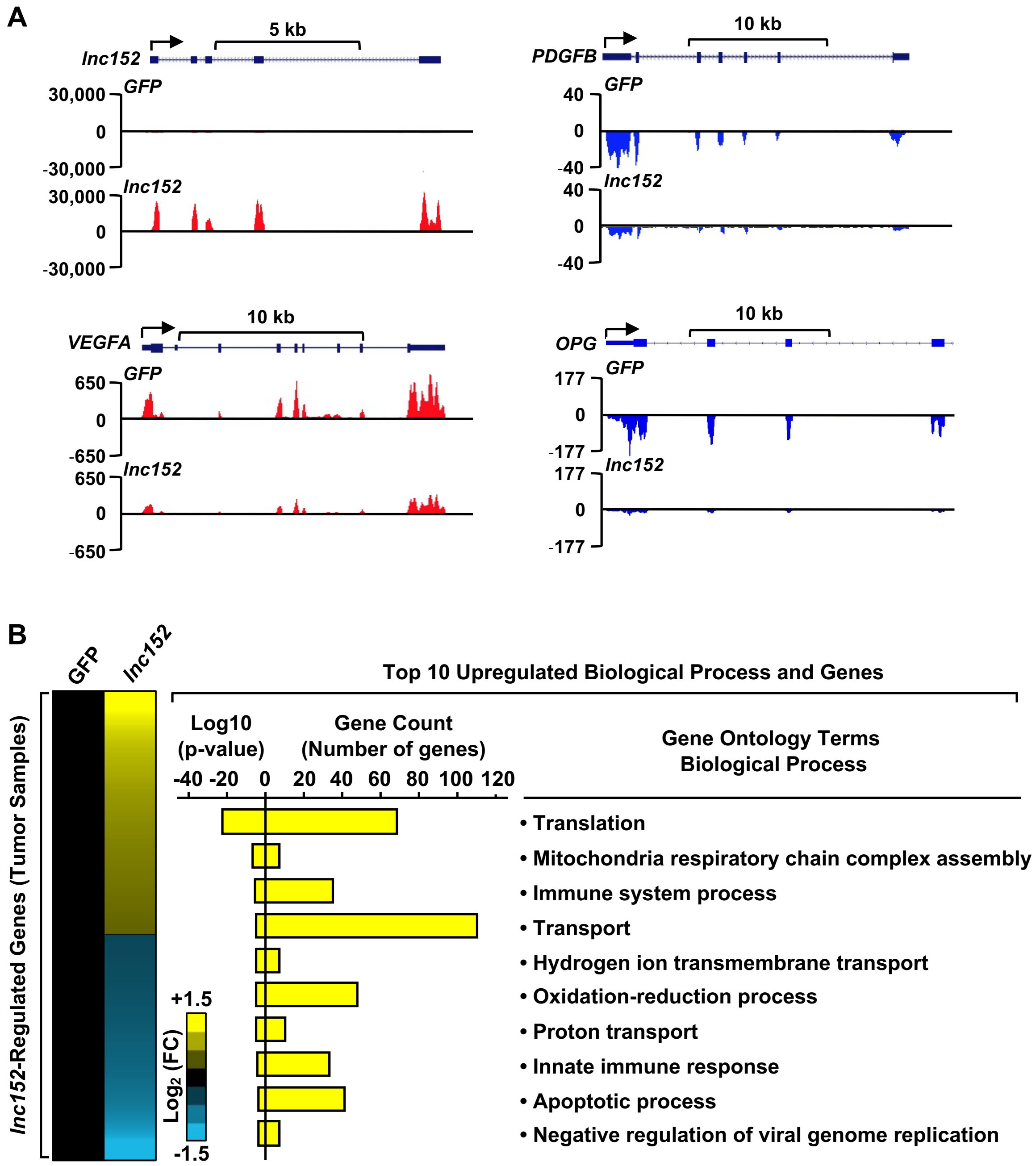

**Figure S4. Ectopic expression of *lnc152* alters gene expression genome-wide.**
**(A)** Genome browser tracks of RNA-seq data showing ectopic expression of *lnc152* with significant down-regulation of selected genes, including *VEGFA*, *PDGFB*, and *OPG* upon ectopic expression of *lnc152*.
**(B)** (*Left*) Heatmap showing RNA-seq data from MDA-MB-231 xenograft tumors engineered for ectopic expression of GFP mRNA or *lnc152*. (*Right*) Gene ontology (GO) analysis showing terms enriched upon ectopic expression of *lnc152* in MDA-MB-231 xenograft tumors relative to expression of GFP mRNA. The GO terms listed indicate the most frequently represented biological processes determined by REViGO analysis.

*[Figure S5 is on the next page]*

**Figure S5. Characterization of MDA-MB-231 cell lines stably expressing luciferase with Dox-inducible expression of exogenous GFP or *lnc152*.**
**(A)** Bioluminescence signals from exogenous GFP or *lnc152* expressing MDA-MB-231 cells after luciferin treatment.
**(B)** Dox-induced ectopic expression of *lnc152* inhibits proliferation of MDA-MB-231 cells stably expressing luciferase compared to ectopic expression of GFP mRNA. Cells were treated daily with 0.5 μg/mL of Dox for six days. Each point represents the mean ± SEM, n = 3. Points marked with asterisks are statistically different from each other (two-tailed Student's t-test, * $p < 0.05$, **** $p < 0.0001$).
**(C)** The expression levels of selected *lnc152*-regulated genes measured upon ectopic expression of *lnc152*, as assessed by RT-qPCR from MDA-MB-231 cells stably expressing luciferase. ND indicates expression was not detectable. Points marked with asterisks are statistically different from each other (two-tailed Student's t-test, * $p < 0.05$, ** $p < 0.01$, **** $p < 0.0001$).
**(D)** Dox-induced ectopic expression of *lnc152* prevents human umbilical vein endothelial cell (HUVEC) tube formation on Matrigel. Morphological appearance of HUVECs grown on Matrigel with conditioned medium (CM) collected from MDA-MB-231 cells expressing Dox-induced GFP or *lnc152*, stained with Calcein-AM (green) and detected by fluorescence microscopy.
**(E)** Quantification of results from the experiments shown in panel D. Image J software with the Angiogenesis plugin was used to detect the number of nodes, number of segments, segment length, and branching length. Each bar represents the mean + SEM, n = 4. Bars marked with asterisks are statistically different from each other (two-tailed Student's t test, * $p < 0.05$, ** $p < 0.01$).
**(F-H)** Dorsal bioluminescence signals from each group immediately after intracardiac tumor cell injection (F) show no statistically significant differences in Luc activity in mice expressing *lnc152* compared to GFP mRNA (G). The dorsal bioluminescence signal from each group immediately after intracardiac tumor cell injection does not correlate with the final outcome (H).

**Figure S5.**

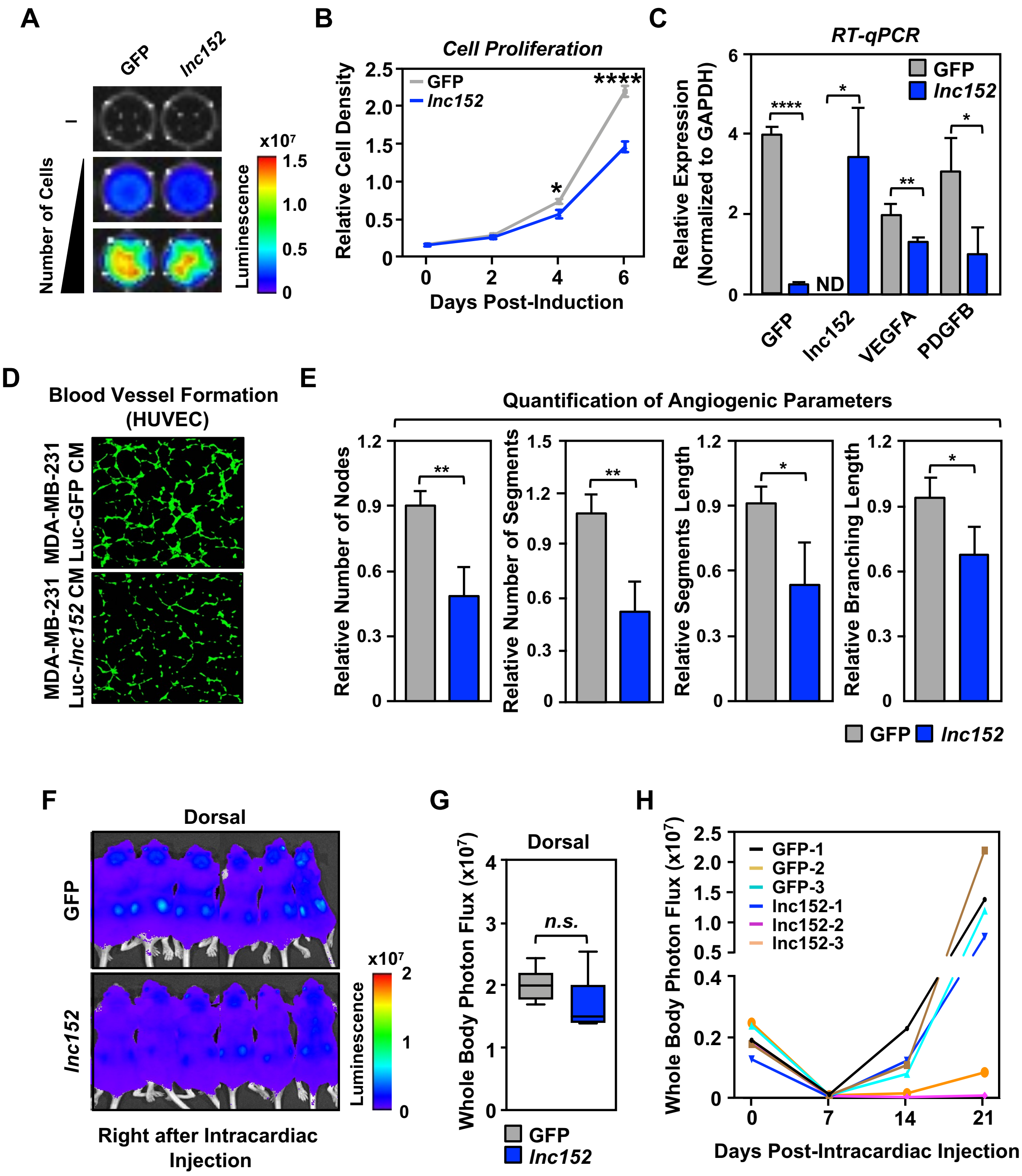
A
GFP
lnc152
–
Number of Cells
x10⁷
1.5
1.0
0.5
0
Luminescence
B
Cell Proliferation
GFP
lnc152
Relative Cell Density
Days Post-Induction
C
RT-qPCR
Relative Expression (Normalized to GAPDH)
ND
GFP
lnc152
VEGFA
PDGFB
D
Blood Vessel Formation (HUVEC)
MDA-MB-231 Luc-GFP CM
MDA-MB-231 Luc-lnc152 CM
E
Quantification of Angiogenic Parameters
Relative Number of Nodes
Relative Number of Segments
Relative Segments Length
Relative Branching Length
GFP
lnc152
F
Dorsal
GFP
lnc152
x10⁷
Luminescence
Right after Intracardiac Injection
G
Dorsal
Whole Body Photon Flux (x10⁷)
n.s.
GFP
lnc152
H
Whole Body Photon Flux (x10⁷)
GFP-1
GFP-2
GFP-3
lnc152-1
lnc152-2
lnc152-3
Days Post-Intracardiac Injection

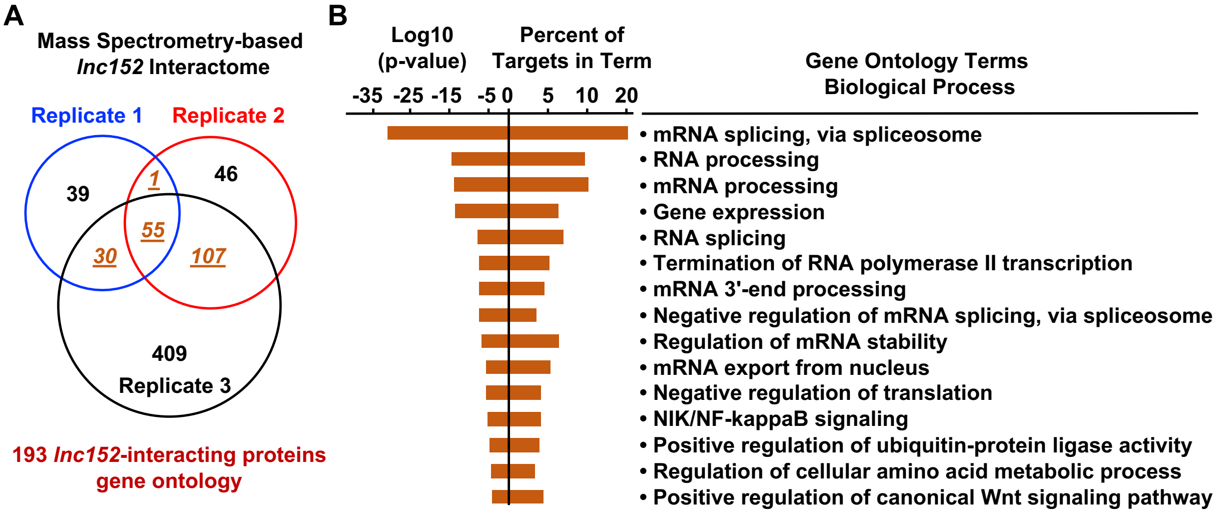

**Figure S6. Analysis of the *lnc152* protein interactome by mass spectrometry.**
**(A)** A diagram depicting the *lnc152*-interactome from three independent RNA pull-down assays. **(B)** Gene ontology (GO) analysis of 193 *lnc152*-interacting proteins identified in at least two independent RNA pull-down assays (highlighted in orange in panel A). The GO terms listed in each cluster indicate the most frequently represented biological processes determined by REViGO analysis.

**A**

**RBM47 Data Set**

Lower RBM47 expression in brain metastatic derivatives of MDA-MB-231

↓

Reintroduction of $RBM47_{WT}$ and $RBM47_{Mutant}$ in 231 derivatives

↓

RNA-seq

↓

RBM47-up and down genes (Supp File 2)

*(Vanharabta et al., 2014)*

***lnc152* Data Set**

Generation of inducible ectopic expression of *lnc152* in MDA-MB-231

↓

Mouse xenograft/ RNA isolation from tumor tissue

↓

RNA-seq

↓

*lnc152*-up and down genes

**B**

**Percent Overlap Between RBM47 and lnc152 Datasets**

| Number of upregulated genes in RBM47 dataset | Upregulated genes common in *lnc152* dataset | Percent Overlap |
|---|---|---|
| 102 | 31 | 30.4% (31/102) |
| **Number of downregulated genes in RBM47 dataset** | **Downregulated genes common in *lnc152* dataset** | **Percent Overlap** |
| 92 | 48 | 48.3% (48/92) |

**C**

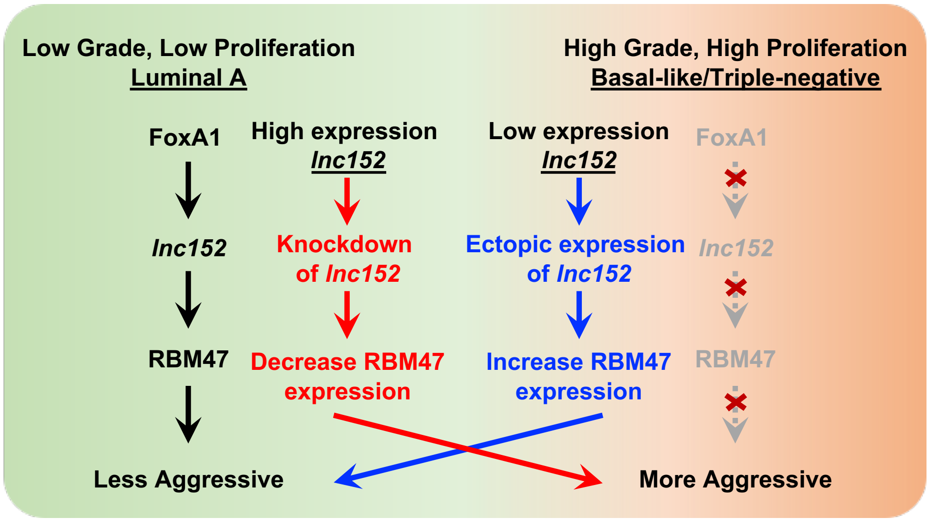

**Figure S7.** ***lnc152*-regulated genes partially overlap with RBM47-regulated genes.**
**(A)** Overview of the pipeline used to analyze the overlap between *lnc152*-regulated genes and RBM47-regulated genes.
**(B)** Percent overlap between *lnc152*-regulated genes and RBM47-regulated genes.
**(C)** Model for the functional link between *lnc152* and RBM47 in the control of tumor aggressiveness. See descriptions in the text.

#### 3) SUPPLEMENTARY MATERIALS AND METHODS

##### Cell Culture

Cancer cell lines MCF-7, MDA-MB-231, T-47D, HCC1143, and 293T cells were purchased from the American Type Cell Culture (ATCC). Primary human mammary epithelial cells (HMEC), and human umbilical vein endothelial cells (HUVEC) were also purchased from the ATCC. Luciferase-expressing MDA-MB-231 cells (1) were kindly provided by Dr. Srinivas Malladi. MCF-7 cells were maintained in Minimum Essential Medium Eagle (Sigma-Aldrich, M1018) supplemented with 5% calf serum and 1% penicillin/streptomycin. MDA-MB-231, T-74D, and HCC1143 cells were maintained in RPMI (Sigma-Aldrich, R8758) supplemented with 10% fetal bovine serum and 1% penicillin/streptomycin. 293T cells were maintained in DMEM (Sigma-Aldrich, D5796) supplemented with 10% fetal bovine serum and 1% penicillin/streptomycin. HMEC were maintained in Mammary Epithelial Cell Basal Medium (ATCC® PCS-600-030™) supplemented with Mammary Epithelial Cell Growth Kit (ATCC® PCS-600-040™). HUVEC were maintained in Vascular Cell Basal Medium (ATCC® PCS-100-030™) supplemented with Endothelial Cell Growth Kit-BBE (ATCC® PCS-100-040™). Luciferase-expressing MDA-MB-231 cells were maintained in DMEM supplemented with 10% fetal bovine serum, 1% penicillin/streptomycin, and 0.5 μg/mL puromycin (Sigma-Aldrich, P9620). Fresh cell stocks were regularly replenished from the original stocks, verified for cell type identity using the GenePrint 24 system (Promega, B1870), and confirmed as mycoplasma-free every three months using a commercial testing kit.

##### Cell Treatments

For doxycycline (Dox; Sigma-Aldrich, D9891) induction, cells were treated with 1 μg/mL of Dox for 24-48 hours. For cell growth assays, cells were pretreated with 1 μg/mL of Dox for 24 hours, re-seeded into 6-well plates, and grown with daily changes of Dox-containing medium (0.5μg/mL) for the indicated times before collection.

##### Antibodies

The antibodies used were as follows: rabbit polyclonal PECAM antibody (Proteintech, 10528-1-AP), rabbit polyclonal RBM47 antibody (Invitrogen, PA5-5432), rabbit polyclonal β-actin antibody (Cell Signaling, #4967), rabbit polyclonal β-tubulin antibody (Invitrogen, PA5-5432), mouse monoclonal FLAG antibody (Sigma-Aldrich, F3165), mouse monoclonal His-probe antibody (Millipore, MABE343).

##### Molecular Cloning to Generate Knockdown and Expression Vectors

***Human cDNA pools.*** cDNA pools were prepared by extraction of total RNA from MCF-7 cells using RNeasy Plus Mini Kit (Qiagen, 74136), followed by reverse transcription using SuperScript III Reverse Transcriptase (Invitrogen, 18080093) with random hexamer primers (Roche, 11034731001) according to the manufacturer's instructions. The cDNA pools were used to amplify *lnc152* and RBM47 cDNAs for subsequent cloning, as described below.

***In vitro transcription vectors.*** A cDNA encoding human *lnc152* was amplified from a cDNA pool by PCR using the primer sets listed below and cloned into *EcoRI*- and *XbaI*-digested pcDNA3 (Thermo Fisher Scientific) downstream of the T7 promoter. A cDNA encoding GFP was amplified from the pcDNA3-GFP expression vector by PCR using the primer sets listed below and cloned into *EcoRI*- and *XbaI*-digested pcDNA3.

*lnc152* Forward: 5’-CGGAATTCGCATGGCCGAATACTGTGTTTTTATC-3’
*lnc152* Reverse: 5’-GCTCTAGAATTTGTATTCACCTTTAATAATGATATGC-3’
GFP Forward: 5’-CGGAATTCTCAGCCACCCGCCACTGCA-3’
GFP Reverse: 5’-GCTCTAGAGCTGTTCCCATGCTTTCGG-3’

***Bacterial expression vectors***. The human RBM47 cDNA was amplified by PCR from a cDNA pool (see above) using the primers listed below, and then cloned into *NotI*- and *XhoI*-digested pET19b (Novagen) bacterial expression vectors to generate a 6xHis-tagged human RBM47 construct.
- *RBM47* Forward: 5’-CGGCGGCCGCACCGGGAAAACTCCGTAGTGACGCT 3’
- *RBM47* Reverse: 5’-GCCTCGAGTTATTGACCAAATGCTTTACTGAAACTCCGC 3’

***Lentiviral expression vectors***. To generate lentiviral vectors for expression of *lnc152*, RBM47, and GFP in MDA-MB-231 cells, the *lnc152*, RBM47, and GFP cDNAs described above were cloned into pINDUCER20 (Addgene, plasmid no. 44012) using a Gibson Assembly kit (NEB, E2621).

**Purification of RBM47 Protein Expressed in *E. coli*.**

6xHis-tagged human RBM47 was expressed in *E. coli* Rosetta (DE3) using the pET19b-based bacterial expression vector described above. The transformed bacteria were grown in LB containing ampicillin and chloramphenicol at 37 °C until the OD595 reached ~ 0.7. Recombinant protein expression was induced by the addition of 0.5 mM IPTG for 14 h at 20 °C. The cells were collected by centrifugation, and the cell pellets were flash-frozen in liquid $N_2$.

The frozen cell pellets were thawed on ice for 1h and lysed by Dounce homogenization (Wheaton) and sonication in His Lysis Buffer (50 mM Tris-HCl pH 7.5, 0.5 M NaCl, 0.1 mM EDTA, 0.1% NP-40, 10% glycerol, 10 mM imidazole, 1 mM PMSF, and 2 mM β-mercaptoethanol). The lysates were clarified by centrifugation at 15,000 rpm using an SS34 rotor (Sorvall) at 4°C for 30 minutes. The supernatant was incubated with 1 mL of Ni-NTA resin equilibrated in Ni-NTA Equilibration Buffer (10 mM Tris-HCl pH 7.5, 0.5 M NaCl, 0.1% NP-40, 10% glycerol, 10 mM imidazole, and 1 mM β-mercaptoethanol) at 4°C for 2 hours with gentle mixing. The resin was collected by centrifugation at 4°C for 10 minutes at 1,000 x *g*, and the supernatant was removed. The resin was washed twice with His Wash Buffer #1 (50 mM Tris pH 7.9, 150 mM NaCl, 2 mM $MgCl_2$, 0.2 mM EDTA, 15 % glycerol, 0.01% NP-40, 10 mM imidazole, 0.2 mM β-mercaptoethanol, 1 mM PMSF, 1 µM aprotinin, 100 µM leupeptin), twice with His Wash Buffer #2 (50 mM Tris pH 7.9, 1 M NaCl, 2 mM $MgCl_2$, 0.2 mM EDTA, 15% glycerol, 0.01% NP-40, 30 mM imidazole, 0.2 mM β-mercaptoethanol, 1 mM PMSF, 1 µM aprotinin, 100 µM leupeptin), and twice with His Wash Buffer #3 (50 mM Tris pH 7.9, 200 mM NaCl, 2 mM $MgCl_2$, 0.2 mM EDTA, 15% glycerol, 0.01% NP-40, 10 mM imidazole, 0.2 mM β-mercaptoethanol, 1 mM PMSF).

The recombinant proteins were eluted using Ni-NTA Elution Buffer (10 mM Tris-HCl pH 7.5, 0.2 M NaCl, 0.1% NP-40, 10% glycerol, 400 mM imidazole, 1 mM PMSF, and 1 mM β-mercaptoethanol). The eluates were collected by centrifugation at 4°C for 10 minutes at 1,000 x *g* and dialyzed in Ni-NTA Dialysis Buffer (10 mM Tris-HCl pH 7.5, 0.2 M NaCl, 10% glycerol, 1 mM PMSF, and 1 mM β-mercaptoethanol). The dialyzed proteins were quantified using a Bradford protein assay (Bio-Rad), aliquoted, flash-frozen in liquid $N_2$, and stored at −80°C. Aliquots of the purified protein were analyzed by SDS-PAGE and staining by Coomassie brilliant blue.

**Generation of Knockdown, Knockout, and Ectopic Expression Cell Lines**

***siRNA-mediated knockdown in MCF-7 cells.*** siRNA oligos targeting human *lnc152* [described previously (2)] were transfected into MCF-7 cells at a final concentration of 10 nM using Lipofectamine RNAiMAX reagent (Invitrogen, 13778150) according to the manufacturer's instructions. After 24 hours or the times indicated, total RNA was isolated using Trizol (Sigma-Aldrich, T9424) and whole cell extracts were prepared as described below to assay for knockdown by RT-qPCR and Western blotting, respectively.

***Dox-inducible ectopic expression in MDA-MB-231 and HCC1143 cells.*** Lentiviruses were generated by transfection of pINDUCER20-based vectors (Addgene, plasmid no. 44012) for Dox-inducible expression of *lnc152*, GFP, and RBM47, along with pCMV-VSV-G, pCMV-GAG-Pol-Rev, and pAdVAntage into 293T cells using Lipofectamine 3000 according to the manufacturer's protocol. After 24 hours, the culture medium was replaced with fresh medium and the cells were maintained for an additional 24 hours. The virus-containing supernatants were collected, filtered through a 0.45 μm syringe filter, and concentrated by using a Lenti-X concentrator (Clontech, 631231). The filtered supernatants were used to infect MDA-MB-231 cells, luciferase-expressing MDA-MB-231 cells, or HCC1143 cells supplemented with 1 μg/mL polybrene (Sigma-Aldrich) to increase the efficiency of viral transduction. Stably transduced cells were isolated under drug selection with 1 μg/mL Geneticin (Life Technologies, 11811031). These engineered cell lines were used for Dox-induced ectopic expression of *lnc152*, GFP, and RBM47 to perform a variety of experiments described herein.

**Cell Proliferation Assays**

Cells expressing *lnc152*, GFP, or RBM47 were plated at a density of 0.2 x $10^5$ cells per well in six well plates and grown with daily changes of Dox-containing medium (0.5 μg/mL) for the indicated times before collection. The cells were washed with PBS, fixed for 10 minutes with 10% formaldehyde at room temperature, and stored in PBS at $4^{\circ}$C until all time points had been collected. The collected cells were stained with a 0.1% crystal violet in 20% methanol solution containing 200 mM phosphoric acid. After washing to remove unincorporated stain, the crystal violet was extracted using 10% glacial acetic acid and the absorbance was read at 595 nm. All growth assays were performed a minimum of three times using independent plantings of cells to ensure reproducibility, and statistically analyzed using Student's unpaired t-test.

**Preparation of Nuclear Extracts and Western Blotting**

***Whole cell lysates from cell lines.*** The cells were washed twice with ice-cold PBS and scraped gently into ice-cold PBS. After the collection by centrifugation, the cell pellet was resuspended in Whole Cell Lysis Buffer (20 mM Tris-HCl pH 7.5, 150 mM NaCl, 1 mM EDTA, 1% NP-40, 1% sodium deoxycholate, 0.1% SDS) containing 1 mM DTT, 1x complete protease inhibitor cocktail (Roche, 11697498001) and phosphatase inhibitors cocktail III (Sigma, P0044-1ML). After a 30 minute incubation on ice with occasional vortexing, the resulting extract was clarified by two rounds of centrifugation at full speed in a microfuge for 10 minutes at 4°C to remove the insoluble material. The supernatant was collected as the soluble whole cell extracts. Protein concentrations for the whole cell extracts were determined using a Bradford protein assay (Bio-Rad). The extracts were aliquoted, flash frozen in liquid $N_2$, and stored at -80°C until used.

***Whole cell lysates from xenograft tumor tissues.*** The frozen xenograft tumor tissues were pulverized using a tissue mill and lysed in Whole Cell Lysis Buffer (20 mM Tris-HCl pH 7.5, 150 mM NaCl, 1 mM EDTA, 1 mM EGTA, 1% NP-40, 1% sodium deoxycholate, 0.1% SDS, 1 mM

DTT, supplemented with protease and phosphatase inhibitors) for extraction of protein as described above.

***Western blotting***. For each extract collected from the indicated conditions, 10 to 20 μg of protein was separated on an 8% to 12% polyacrylamide-SDS gel and transferred to a nitrocellulose membrane. The membranes were blocked for 1 hour at room temperature in Tris-Buffered Saline (TBS) with 0.05% Tween (TBST) containing 5% nonfat dry milk. Primary antibodies PECAM (1:2000), RBM47 (1:2000), FLAG (1:5000), His-probe (1:5000), β-actin (1:5000), or β-tubulin (1:5000) were diluted in 5% nonfat dry milk and incubated with membranes overnight at 4°C with gentle mixing. After extensive washing with TBST, the membranes were incubated with an appropriate HRP-conjugated secondary antibody (Pierce) diluted in 1% nonfat dry milk for 1 hour at room temperature. The membranes were washed extensively with TBS-T before chemiluminescent detection using SuperSignal West Pico substrate (ThermoFisher Scientific) and X-ray film or ChemiDoc system (Bio-Rad).

**RNA Isolation and Reverse Transcription Quantitative PCR (RT-qPCR)**

***RNA isolation and preparation of cDNA samples***. Cells expressing inducible *lnc152*, GFP, or RBM47 were seeded in 6-well plates and treated with 1 μg/mL Dox for 24-48 hours. The cells were collected at the indicated time points and cDNA pools were prepared by extraction of total RNA from cell lines using Trizol (Sigma- Aldrich, T9424), followed by reverse transcription using MMLV reverse transcriptase (Promega, M150B) with random hexamer or oligo(dT) primers (Sigma-Aldrich). The cDNA was treated with 3 units of RNase H (Ambion) for 30 minutes at 37°C and was used for qPCR reactions.

***qPCR***. The cDNA samples were subjected to qPCR using gene-specific primers, as listed below. Briefly, cDNA, 1x SYBR Green PCR master mix, and forward and reverse primers (250 nM) were mixed and subjected to 45 cycles of amplification (95°C for 10 seconds, 60°C for 10 seconds, 72°C for 1 second) following an initial 5 min incubation at 95°C using a LightCycler 480 real-time PCR thermocycler (Roche). Melting curve analyses were performed to ensure that only the targeted amplicon was amplified. Target gene expression was normalized to the expression of human GAPDH or RPL19. Relative expression was determined in comparison to the value of an appropriate control sample. Statistical differences between two groups were determined using the Student's t-test and a statistical significance was defined as $p < 0.05$.

***Primers used for RT-qPCR***. The following primers were used for RT-qPCR.

| | |
|---|---|
| *GFP* forward: | 5'-ACGACGGCAACTACAAGACC-3' |
| *GFP* reverse: | 5'-TTGTACTCCAGCTTGTGCCC-3' |
| *lnc152* forward: | 5'-AGAAATGCCACCGGACATAG-3' |
| *lnc152* reverse: | 5'-CATACTTCTGCTGCGTCCAA-3' |
| *FOXA1* forward: | 5'-GGGGGTTTGTCTGGCATAGC-3' |
| *FOXA1* reverse: | 5'-GCACTGGGGGAAAGGTTGTG-3' |
| *VEGFA* forward: | 5'-GCACCCATGGCAGAAGG-3' |
| *VEGFA* reverse: | 5'-CTCGATTGGATGGCAGTAGCT-3' |
| *PDGFB* forward: | 5'-GGGCAGGGTTATTTAATATGG-3' |
| *PDGFB* reverse: | 5'-AATCAGGCATCGAGACAG-3' |
| *OPG* forward: | 5'-ATAGATGTTACCCTGTGTGAG-3' |
| *OPG* reverse: | 5'-AAGACACTAAGCCAGTTAGG-3' |
| *ETS1* forward: | 5'-CTCAGATATGGAATGTGCAG-3' |
| *ETS1* reverse: | 5'-TGCTGTTCTTTAGTGAAACC-3' |

| | |
|---|---|
| *FGF1* forward: | 5'-TACTCTGAGAAGAAGACACC-3' |
| *FGF1* reverse: | 5'-GCGCTTTCAAGACTAAAGAG-3' |
| *FOXS1* forward: | 5'-GCTCTAGGACCTGAAGAAC-3' |
| *FOXS1* reverse: | 5'-TTCCCTCATCATCACATTGG-3' |
| *KFI14* forward: | 5'-TTCAGAACACCTCTGCAGGA-3' |
| *KFI14* reverse: | 5'-ACTCATGAAGACTACCTGGG-3' |
| *FSCN1* forward: | 5'-GGAGACCGACCAGGAGAC-3' |
| *FSCN1* reverse: | 5'-CATTGGACGCCCTCAGTG-3' |
| *TRIM25* forward: | 5'-AAGAAATCCAAGAAACCTCC-3' |
| *TRIM25* reverse: | 5'-CTTGAGAGATGTTGAGTTCG-3' |
| *PGF* forward: | 5'-AGCTCCTAAAGATCCGTTC-3' |
| *PGF* reverse: | 5'-GACGGTAATAAATACACGAGC-3' |
| *HMOX1* forward: | 5'-CAACAAAGTGCAAGATTCTG-3' |
| *HMOX1*reverse: | 5'-TGCATTCACATGGCATAAAG-3' |
| *MMP2* forward: | 5'-TTCTGGAGATACAATGAGGTG-3' |
| *MMP2* reverse: | 5'-CTTGAAGAAGTAGCTGTGAC-3' |
| *ANTXR1* forward: | 5'-GAGAGTCATTTCAAGTTGTCG-3' |
| *ANTXR1* reverse: | 5'-GAGTCATTGATCTTGAAGCTG-3' |
| *BMP2* forward: | 5'-TCCACCATGAAGAATCTTTG-3' |
| *BMP2* reverse: | 5'-TAATTCGGTGATGGAAACTG-3' |
| *RBM47* forward: | 5'-GAAGATGTTGTAGGTGACTTC-3' |
| *RBM47* reverse: | 5'-CCTGAGCTTTCCTTGTTAAG-3' |
| *GAPDH* forward: | 5'-CCACTCCTCCACCTTTGA-3' |
| *GAPDH* reverse: | 5'-ACCCTGTTGCTGTAGCCA-3' |
| *RPL19* forward: | 5'-ACATCCACAAGCTGAAGGCA-3' |
| *RPL19* reverse: | 5'-TGCGTGCTTCCTTGGTCTTA-3' |
| *ACTB* forward: | 5'-CATGTACGTTGCTATCCAGGC-3' |
| *ACTB* reverse: | 5'-CTCCTTAATGTCACGCACGAT-3' |

**RNA-sequencing**

***Generation of RNA-seq libraries***. Total RNA from seven xenograft tumor tissues of each group was extracted using the RNeasy Plus Kit (Qiagen, 74134) according to the manufacturer's instructions and used to generate strand-specific RNA-seq libraries. The RNA was then enriched for polyA+ RNA using Dynabeads Oligo(dT)25 (Invitrogen, 61002). The polyA+ RNA was then used to generate strand-specific RNA-seq libraries as described previously (3). The RNA-seq libraries were subjected to QC analyses (i.e., number of PCR cycles required to amplify each library, the final library yield, and the size distribution of the final library DNA fragments) and sequenced using an Illumina HiSeq 2500 and NextSeq 500.

***Analysis of RNA-seq data sets***. The raw data were subjected to QC analyses using the FastQC (4). The raw reads were aligned to the human reference genome (hg19/GRCh37) using default parameters in Tophat (v2.0.12) (5). Uniquely mappable reads were converted into bigWig files using BEDTools (6) for visualization in the Integrative Genomics Viewer (7). Transcriptome assembly was performed using cufflinks v.2.2. (8) with default parameters. The transcripts were merged into distinct, non-overlapping sets using cuffmerge, followed by cuffdiff to call the differentially regulated transcripts (8). The significantly ($q < 0.001$) regulated genes were determined by comparing the experimental samples to corresponding untreated control samples to determine the regulated gene sets. The differentially expressed genes identified from the analysis described above were used in a number of subsequent downstream analyses and the data were visualized using a variety of approaches.

***Data visualization and statistics***. Box plot representations were used to quantitatively assess the $\log_2$ fold changes in gene expression in the different experimental conditions compared to matched untreated controls. Box plots were generated using custom scripts in R. Wilcoxon rank sum tests were performed to determine the statistical significance of all comparisons. Line plots were generated using custom scripts in R to represent the trend of the $\log_2$ fold changes in gene expression in the different experimental conditions compared to matched untreated controls. Browser tracks were generated using bigWig files that represented fold change in signal for each condition relative to its input. Browser tracks were visualized using visualization in the Integrative Genomics Viewer (7).

***Gene set enrichment analysis***. Gene ontology (GO) analyses were performed using the DAVID (Database for Annotation, Visualization, and Integrated Discovery) tool (9). DAVID returns clusters of related ontological terms that are ranked according to an enrichment score.

**Mining of Public Databases**

The expression of *lnc152* and RBM47 in normal breast and different subtypes of breast cancer was determined using GEPIA based on the RPKM values in the TCGA dataset (10). GEPIA analyzes tumors and normal samples from the TCGA and the GTEx projects, using a standard processing pipeline and provides differential expression analysis for gene profiling according to cancer subtypes.

**Analysis of ChIP-seq Data**

FoxA1 ChIP-seq libraries from MCF-7 cells (NCBI GEO accession number GSE59530) were generated and analyzed as described previously (11).

**Analysis of RBM47- and *lnc152*-Regulated Genes**

RBM47 up- and down-regulated genes were identified from a dataset generated from brain metastatic derivatives of MDA-MB-231 in which wild-type or mutant RBM47 was introduced (12). The list was then compared to the list of *lnc152* up- and down-regulated genes determined from our dataset. Overlaps of common up- and down-regulated genes were represented as percent overlap genes in the *lnc152* regulated genes in common with the RBM47 dataset.

**In Vitro Transcription to Generate GFP mRNA and *lnc152***

To generate GFP mRNA or *lnc152* in vitro, *XbaI*-linearized plasmid DNA containing cDNAs for the genes of interest (pcDNA3-GFP or pcDNA-*lnc152*) were purified on an agarose gel and transcribed with T7 RNA polymerase (Promega, P207B) according to the manufacturer's protocol. One μg of the linearized plasmid DNA was incubated with T7 RNA polymerase and NTPs (10 mM each of ATP, CTP, GTP, and UTP) at 37°C for 2-3 hours. The template DNA was then removed by treatment with DNase (Promega, M6101) at 37°C for 15 minutes. The reaction was stopped by the addition of 2 μL of 0.2 M EDTA (pH 8.0). The in vitro transcribed RNAs were purified using an RNeasy Plus Kit and the sizes of the transcripts were confirmed using an RNA BioAnalyzer. To generate biotin-labeled RNAs for pull down assays, an NTP labeling mixture containing biotin-UTP (10 mM each ATP, CTP, and GTP, 6.5 mM UTP, 3.5 mM biotin-UTP; Sigma, 11685597910) was used for in vitro transcription.

**In Vitro RNA Pull Down Assays Combined with LC-MS/MS Analysis**

***In vitro RNA pull down***. GFP mRNA or *lnc152* were transcribed and labeled with biotin in vitro as described above, and then pre-folded in Folding Buffer (10 mM Tris-HCl pH 7.4, 100 mM NaCl, 1 mM EDTA) by incubating for 2 minutes at 70˚C, 5 minutes at 50˚C, 15 minutes at 37˚C, and 15 minutes at room temperature. After incubation, 50 mM KCl and 10 mM $MgCl_2$ (final concentrations) were added, and the RNA was incubated on ice for 10 minutes and stored on ice until used. Whole cell extract from MCF-7 cells, prepared as described above, was incubated with 5 nM of folded RNA in Binding Buffer (20 mM HEPES pH 7.9, 150 mM NaCl, 50 mM KCl, 10 mM $MgCl_2$, 0.2 mM $ZnCl_2$, 0.1% BSA, 0.2% NP-40, 5% glycerol, 15 mM β-mercaptoethanol, 0.2 mg/mL yeast tRNA, 1 mg/mL BSA, 100 U/mL SUPERaseIn) at room temperature for 1 hour, followed by incubation with equilibrated Streptavidin-agarose beads (ThermoFisher Scientific, 20377) at room temperature for 30 minutes. The beads containing the protein-RNA complex were washed four times in Binding Buffer containing an additional 200 mM of NaCl. The reactions were stopped by the addition of 4x SDS-PAGE Loading Buffer, followed by heating to 100˚C for 10 minutes. Aliquots were analyzed by SDS-PAGE with silver staining to visualize interacting proteins.

***LC-MS/MS analysis.*** In order to detect the RNA-bound proteins by mass spectrometry, the samples were resolved on a 4-12 % acrylamide-SDS gel (Invitrogen, NW04120BOX) and visualized by Coomassie blue staining. Gel slices containing the RNA-bound proteins were excised and transferred to a microfuge tube. Following reduction and alkylation with DTT and iodoacetamide (Sigma-Aldrich, A3221), respectively, the RNA-bound proteins in the gel slice were digested overnight with trypsin (Promega, V5111). The samples were then subjected to solid-phase extraction cleanup with an Oasis HLB plate (Waters) and the resulting samples were injected onto an Orbitrap Fusion Lumos mass spectrometer (Thermo Electron) coupled to an Ultimate 3000 RSLC-Nano liquid chromatography system (Dionex).

The samples were injected onto a 75 µm i.d., 50-cm long EasySpray column (Thermo) and eluted with a gradient from 1-28% Buffer B over 60 minutes. Buffer A contained 2% (v/v) ACN and 0.1% formic acid in water, and Buffer B contained 80% (v/v) ACN, 10% (v/v) trifluoroethanol, and 0.1% formic acid in water. The mass spectrometer operated in positive ion mode with a source voltage of 1.5-2.4 kV and an ion transfer tube temperature of 275°C. MS scans were acquired at 120,000 resolution in the Orbitrap and up to 10 MS/MS spectra were obtained in the ion trap for each full spectrum acquired using higher-energy collisional dissociation (HCD) for ions with charges 2-7. Dynamic exclusion was set for 25 s after an ion was selected for fragmentation.

Raw MS data files were converted to a peak list format and analyzed using the central proteomics facilities pipeline (CPFP), version 2.0.3. Peptide identification was performed using the X!Tandem and open MS search algorithm (OMSSA) search engines against the human protein database from Uniprot, with common contaminants and reversed decoy sequences appended. Fragment and precursor tolerances of 10 ppm and 0.5 Da were specified, and three missed cleavages were allowed.

***Detection of lnc152-RBM47 interactions.*** The eluted protein-RNA complexes from whole cell extracts described above were subjected to Western blot analyses. For detection of *lnc152*-RBM47 interactions using recombinant proteins, 50 ng of purified His-tagged recombinant RBM47 protein was incubated with 5 nM of folded RNAs in Binding Buffer (20 mM HEPES pH 7.9, 150 mM NaCl, 50 mM KCl, 10 mM $MgCl_2$, 0.2 mM $ZnCl_2$, 0.1% BSA, 0.2% NP-40, 5% glycerol, 15 mM β-mercaptoethanol, 0.2 mg/mL yeast tRNA, 1 mg/mL BSA, 100 U/mL

SUPERaseIn) at room temperature for 1 hour, followed by incubation with equilibrated Streptavidin-agarose beads (ThermoFisher Scientific, 20377) at room temperature for 30 minutes. The beads containing the protein-RNA complex were washed five times in Binding Buffer containing an additional 200 mM of NaCl. The reactions were stopped by the addition of 4x SDS-PAGE Loading Buffer, followed by heating to 100°C for 10 minutes. The samples were then analyzed by Western blotting as indicated.

**Cell Migration and Invasion Assays**

Boyden chamber assays were used to determine the migration and invasive capacity of cells using the following protocol. Cells were pretreated with 1 μg/mL of Dox for 48 hours, then detached and rinsed two times with serum-free medium. After centrifugation, the cell pellet was resuspended in serum-free medium at a cell density adjusted to $2 \times 10^5$ cells/mL. For migration assays, 200 μL of cell suspension was added to trans-well chambers (Corning, 353097) and 750 μL of RPMI1640 medium supplemented with 10% FBS (serving as the chemo-attractant) was added to the outer chamber. For invasion assays, 200 μL of cell suspension was added to trans-well chambers with Matrigel (Corning, 354480). Trans-well chambers with Matrigel were also manually prepared by incubating a mixture of BD Matrigel Basement Membrane Matrix (BD BioScience) and PBS at least 2 hours at 37°C to allow polymerization. After 24 hours incubation at 37°C, the insert chamber was removed and non-migrating cells on the upper surface of the insert chamber were removed with a cotton swab. Afterward, the cells that migrated into the lower side of the membrane were fixed and stained with 0.5% crystal violet in 20% methanol solution for 15 minutes, washed with water, and air-dried. Images were acquired with an upright microscope, and the transmembrane cells were counted in a number of random fields.

**In Vitro Endothelial Tube Formation Assay**

***Preparation of conditioned medium (CM).*** MDA-MB-231 cells were pretreated with 1 μg/mL of Dox for 24 hours, washed three times with serum-free medium, and incubated for 24 hours with serum-free medium in the presence of 1 μg/mL of Dox. The conditioned medium was collected and concentrated using Centricon 10 (Millipore) centrifuge filter unit, aliquoted, flash-frozen in liquid $N_2$, and stored at −80 °C.

***Endothelial tube formation assay.*** 30 μL of Matrigel Basement Membrane Matrix were added onto 24-well plate and incubated at least 1 hour at 37°C to solidify Matrigel. HUVECs were then resuspended at a density of $0.2 \times 10^5$ cells per well by gentle mixing by pipetting in conditioned medium of MDA-MB-231 cells expressing GFP mRNA or *lnc152*. Resuspended HUVECs were seeded onto Matrigel Basement Membrane Matrix coated 24-well plate and incubated at 37°C for 4-12 hours. Capillary-like tubes were then stained with cell-permeable dye Calcein AM (ThermoFisher, C1430) and imaged by fluorescence microscopy.

***Quantification of tube networks.*** The total number of nodes, number of segments, total segment length, and branching were analyzed by the Angiogenesis Analyzer plugin in ImageJ. The normalized values were averaged for three biological replicates with three to four fields in each replicate.

**Indirect Immunofluorescence**

***Staining.*** Xenograft tissues were processed for paraffin sectioning using standard protocols. For immunofluorescent staining, 5 μM paraffin sections were deparaffinized in xylene and rehydrated sequentially in 100%, 95%, 80%, 70%, and 50% ethanol prepared in $ddH_2O$.

Antigen retrieval was performed by boiling the sections in 10 mM citrate pH 6.0 for 10 minutes. After cooling to room temperature for 1 hour, the sections were incubated in 3% $H_2O_2$ in methanol for 12 minutes. After washing with PBS, the sections were then blocked with 5% normal goat serum (Thermo Fisher Scientific, 50062Z) in PBS at room temperature for 1 hour and then incubated with primary PECAM (1:300) antibodies overnight at 4°C in PBS containing 5% normal goat serum. After washing with PBS, the sections were incubated with Alexa fluor 488-conjugated goat anti-rabbit IgG (1:400; ThermoFisher, A-11008), in PBS containing 5% normal goat serum at room temperature for 1 hour. After washing with PBS, the sections and coverslips were mounted with Anti-Fade mounting medium with DAPI (Vector Laboratories, H-1200). Immunofluorescence staining was imaged using a Zeiss LSM880 confocal microscope purchased with a shared instrumentation grant from the NIH (S10OD021684 to Katherine Luby-Phelps).

***Confocal imaging.*** Immunofluorescent staining was imaged using a Zeiss LSM880 confocal microscope (Live Cell Imaging Facility, UT Southwestern Medical Center) purchased with a shared instrumentation grant from the NIH (S10 OD021684 to Katherine Luby-Phelps). A number of 20x magnification fields were photographed from each section from two animals in each group.

**Experiments with Mice**

All animal experiments were performed in compliance with the Institutional Animal Care and Use Committee (IACUC) at the UT Southwestern Medical Center. Female athymic nude (*Foxn1$^{nu}$*; Envigo #069) mice at 6-8 weeks of age were used. We used female mice because mammary cancers occur primarily in females. In addition, the human cancer cell lines that we used for xenografts were obtained from females.

***Xenograft experiments.*** To establish breast cancer xenografts, MDA-MB-231 cells engineered for Dox-inducible expression of GFP mRNA or *lnc152* (5 x $10^6$ in 100 µL) were injected subcutaneously into the flank of the mice in a 1:1 ratio PBS and Matrigel (Fisher, CB 40230). Ten days post-tumor cell injection, the mice were placed on a Dox-containing diet (625 mg/kg; Envigo). Seven mice from each group carrying 100 $mm^3$ subcutaneous tumors were selected to maintain for 3 weeks. Tumor growth was measured using electronic calipers approximately every 3-4 days. Tumor volumes were calculated using a modified ellipsoid formula: tumor volume = ½ (length x width$^2$). At the end of the experiment, the mice were euthanized to collect the xenograft tissue and measure xenograft tissue weight. The tissue was cut into several small pieces and separate portions were either snap-frozen in liquid nitrogen or fixed using 10% formalin. Statistical analyses for the xenograft-based experiments were performed using GraphPad Prism 7 (Student's unpaired t-test).

***Metastasis experiments***. MDA-MB-231 cells harboring a luciferase reporter were engineered for Dox-inducible expression of GFP mRNA or *lnc152.* Briefly, 2 x $10^6$ cells/ml were collected at 75-90% confluence with 0.05% trypsin/EDTA and resuspended in ice-cold PBS (without $Ca^{2+}$ and $Mg^{2+}$). The cell suspension was mixed gently and 100 µL was injected intracardially into the arterial circulation of 5 weeks old female athymic nude (*Foxn1$^{nu}$*) mice (Envigo #069) to mimic hematogenous dissemination (13). Successful injection was confirmed by the detection of luciferase signal in the whole mouse.

***Whole-body bioluminescent imaging***. Mice were monitored weekly for metastatic outgrowth by bioluminescence imaging. D-luciferin (150 mg/kg) was injected retro-orbitally and the mice were imaged using IVIS Spectrum with Living Image 4.4 software (PerkinElmer).

**Quantification and Statistical Analysis**

All sequencing-based genomic experiments were performed a minimum of two times with independent biological samples. Statistical analyses for the genomic experiments were performed using standard genomic statistical tests as described above. All gene specific qPCR-based experiments were performed a minimum of three times with independent biological samples. Statistical analyses for the qPCR-based experiments were performed using GraphPad Prism 7. All tests and p-values are provided in the corresponding figures or figure legends. In all figures, the p values are shown as: *, $p<0.05$; **, $p<0.01$; ***, $p<0.001$, ****, $p<0.0001$.

**Data and Code Availability**

***Genomic datasets.*** The new RNA-seq data generated for this study can be accessed from the NCBI's Gene Expression Omnibus (GEO) repository (http://www.ncbi.nlm.nih.gov/geo/) using the superseries accession number GSE193634.

***Custom scripts.*** Custom R scripts for genomic data analyses are available from the corresponding author.

***Mass spectrometry datasets.*** The new mass spectrometry datasets generated for this study can be accessed from MassIVE using the accession number MSV000088680 and also are available as supplementary data provided with this manuscript (Supplementary Table S3).
